# Gut microbiota variation across sympatric stingless bee species and honey bees in the Neotropics

**DOI:** 10.1101/2025.11.29.691224

**Authors:** Karen Luisa Haag, Luiza Quadro Stein, Carlos Gustavo Nunes da Silva, Florent Mazel, Aiswarya Prasad, Philipp Engel

## Abstract

Stingless bees (Meliponini) are ecologically and culturally important pollinators with a long tradition of human management in the Neotropics. Yet, little is known about how their gut microbiota vary across geographic regions or whether microbial exchange occurs with managed honey bees (*Apis mellifera*), which are often kept in close proximity. Using full-length 16S rRNA gene sequencing of individual bees sampled from 167 colonies, we characterized gut microbial community structure through a hierarchical, taxonomic and phylogenetic comparative framework, contrasting the microbiota of *Melipona quadrifasciata* and *Melipona mondury* with that of *Apis mellifera* across their shared geographic range in Brazil. The core microbiota of *Melipona* was dominated by *Lactobacillus*, *Bifidobacterium*, *Apilactobacillus*, *Bombella*, and *Floricoccus*, and showed inverse variation in relative abundance with lower-prevalence bacterial taxa. Although the core microbiota of the two stingless bee species overlapped only partially with that of *Apis mellifera*, they exhibited comparable alpha-diversity and beta-diversity dispersion, indicating broadly similar community assembly processes and dynamics. Nevertheless, we found that 6% of all amplicon sequence variants (ASVs) were shared between hosts, encompassing nearly all canonical honey bee “core” symbionts, indicating frequent spillover. Remarkably, several ASVs of *Snodgrassella*, a genus typically rare in stingless bees, reached high abundance in several *M. quadrifasciata* individuals and formed a deeply divergent clade (∼96% 16S rRNA gene identity to *S. alvi*). These patterns are consistent with the hypothesis that human-mediated management practices, such as mixed apiaries and artificial feeding, create opportunities for microbial exchange between native and non-native bees. Together, our findings indicate that stingless bee gut microbiomes are compositionally stable yet ecologically permeable, shaped by both long-term host specificity and recent anthropogenic contact.

## Introduction

The gut microbiota of social bees, such as bumble bees, honey bees, and stingless bees, is taxonomically simple and becomes established soon after adult emergence [1, 2]. These symbionts contribute to digestion of pollen- and plant-derived polysaccharides [3, 4], detoxification of secondary plant metabolites [5], host learning and memory [6, 7], and protection against pathogens [8–10]. Because their microbiota are dominated by a small number of host-specific lineages, social bees have become valuable models for studying the evolution, maintenance, and host interactions of animal-associated microbial communities [11]. Yet our understanding of these symbioses is shaped overwhelmingly by the Western honey bee (*Apis mellifera*), the most widely studied and heavily managed bee species. The honeybee gut microbiota is remarkably consistent: five to nine dominant genera, including *Gilliamella, Snodgrassella, Bombilactobacillus, Lactobacillus, Bifidobacterium, Frischella, Bombella, Bartonella* and *Commensalibacter*, account for most bacterial cells in the gut of an adult worker bee [12]. Most of these bacteria have been cultured, and studies with gnotobiotic honeybees have started to reveal their relative contribution to interactions with the host [7, 13, 14]. Different honey bee species harbor gut communities that are similar in composition but host-specific - that is, they share several bacterial genera, but these are mostly represented by distinct species and strains [15, 16]. Some of the same bacterial genera are also present in the gut microbiota of bumble bees and stingless bees, pointing to a shared evolutionary history. However, much less is known about the gut microbiota of these groups of social bees. In particular, knowledge about stingless bees (Meliponini), a taxonomically diverse and ecologically critical clade of tropical pollinators, remains limited. Only recently have studies begun to describe their gut microbiota composition, and most reports are geographically limited [15, 17–20]. We still know little about how variable stingless bee gut communities are across the hosts’ broad geographic ranges, whether they display the same strong host specificity seen in *A. mellifera*, and which ecological or anthropogenic factors might drive community turnover.

Humans have interacted with social bees for millennia. Honey exploitation dates back to the Neolithic [21, 22], and stingless bees were managed by pre-Columbian civilizations long before the worldwide spread of *A. mellifera* by European settlers [23]. Today, meliponicultures and apicultures often occur side by side. Managed colonies of honey bees and stingless bees are frequently kept in close proximity, transported across regions, and manipulated for honey production. Artificial feeding, including supplementing stingless bees with *A. mellifera* honey, is common in times of resource scarcity. These practices create ecological contact zones where gut microbes from one host can encounter and potentially colonize another. Evidence for such microbiota spillover in bees is scarce but growing. Recent evidence from honey bees suggest that some bacteria may be shared among closely related hosts [16]. Some floral- and environment-associated bacteria appear in multiple bee hosts and colony environments [17, 19, 24], and there are scattered reports of *Snodgrassella* or *Lactobacillus* strains occurring outside their typical hosts [18, 25, 26]. However, to the best of our knowledge no broad-scale, strain-level test has addressed whether gut symbionts from managed honey bees infiltrate native stingless bees or vice versa.

High-throughput sequencing of full-length 16S rRNA genes using PacBio now enables the detection of bacterial community composition at the sub-species or strain level, allowing researchers to characterize fine-scale diversity patterns within and across host species. Here, we implement these advances to investigate the gut microbiota of two widespread stingless bees, *Melipona quadrifasciata* and *M. mondury*, sampled alongside sympatric *A. mellifera* colonies across Brazil. Our primary goals were (i) to quantify within- and between-host variation in the gut microbiota of these two important stingless bee species across a broad geographic range, (ii) to test whether these communities are more environmentally assembled or variable than those of honey bees, and (iii) to assess the extent and direction of cross-species bacterial sharing and potential spillover, particularly in human-managed settings where honey bees and stingless bees coexist in close proximity.

## Material and Methods

### Sampling

All bees were sampled between October 2023 and February 2024 and kept in 100% ethanol at -20°C until use (metadata available in **Supplementary File 1**). Sampling was authorized by SISBIO/ICMBio/MMA through the license #89616-1 to KLH. *M. quadrifasciata* subspecies (*M.q. quadrifasciata* and *M.q. anthidioidis*) were recognized by their differences in abdomen coloration and mitochondrial *cox1* gene haplotypes ([27]; see below).

### DNA extraction and amplicon sequencing

Abdomens were removed from bee bodies using clean scalpels. The abdomen of one single bee per colony was used for DNA extraction using the Vazyme FastPure Blood/Cell/Tissue/Bacteria DNA Isolation Mini Kit (Nanjing, China). Briefly, individual abdomens were washed twice in 1X PBS, grinded with a small sterile plastic pestle in a 1.5 ml tube containing 1X G2 buffer (Qiagen, Hilden, Germany) with 20 mg/ml lysozyme, and then incubated at 37°C for at least 1h. DNA extraction was performed according to the manufacturer protocol. DNA quality and yield were evaluated on a Qubit fluorometer (Thermo Fisher Scientific, Waltham, Massachusetts).

For bacterial amplicon sequencing, the full length of the 16S rRNA gene was amplified from each DNA sample with the KAPA HiFi HotStart polymerase (Roche, Basel, Switzerland), using 1 ng of template DNA, barcoded primers GCATC/barcode/AGRGTTYGATYMTGGCTCAG and GCATC/barcode/RGYTACCTTGTTACGACTT, and following the manufacturer instructions. PCR products were pooled, and sequencing libraries were prepared with the SMRTbell Express TPK 2.0 Library Construction system (Pacific Biosciences, Menlo Park, California) using the pooled 16S rRNA gene PCR amplicons. Library preparation, sequencing, data quality control and demultiplexing were performed at the Lausanne Genome Technologies Facility (UNIL, Lausanne, Switzerland).

### *M. quadrifasciata* subspecies identification

The main difference between *M. quadrifasciata* subspecies refers to the yellow metasomal stripes, which are continuous in *M. q. quadrifasciata* and discontinuous in *M. q. anthidioides* [28], but this feature is often ambiguous. Therefore, in addition to recording the coloration pattern and the information provided by beekeepers upon sampling, barcoding of mitochondrial *cox1* gene was used to help classifying *M. quadrifasciata* samples in each of the two recognized subspecies. To this end, a DNA aliquot was used to amplify the *cox1* gene with primers BarbeeF [29] and MtD9 [30], generating ∼619 bp amplicons. Amplicons were purified with the Exo-CIP Rapid PCR Cleanup Kit (New England Biolabs, Ipswich, Massachusetts) according to the manufacturer instructions and then sequenced with the Sanger method by Microsynth (Balgach, Switzerland). For the analysis, the Sanger sequences were assessed for quality and aligned using the Muscle algorithm [31] within Geneious Prime 2025.2.1 (Biomatters, Auckland, New Zealand) and imported into PopART (http://popart.otago.ac.nz) for haplotype identification. A network was built using the TCS algorithm [32]. Subspecies were assigned to separate groups of haplotypes within the network (**Fig. 1D**), and validated by the information collected at the sampling sites.

**Figure 1.**
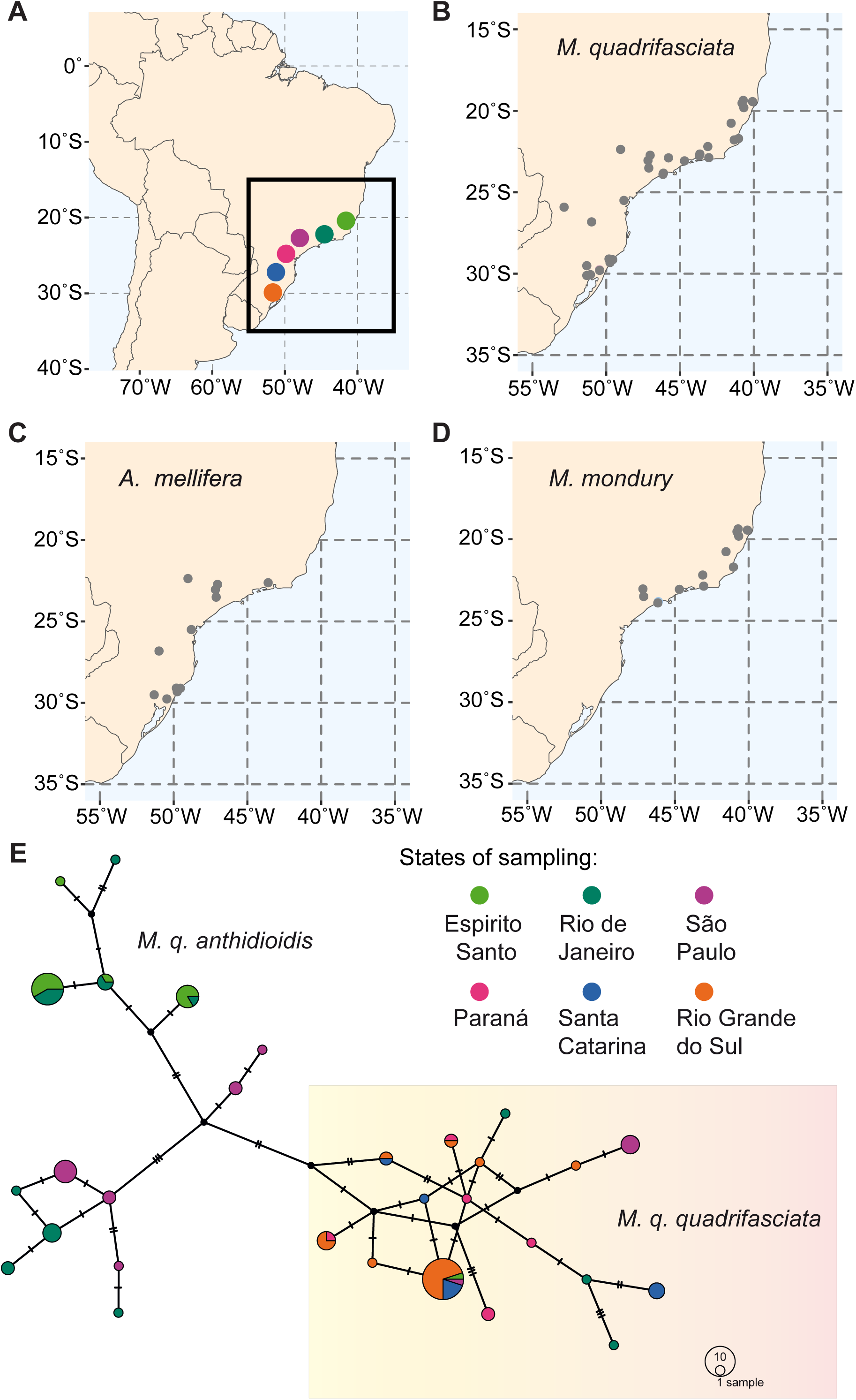
Origin of bees analyzed in our study. **A.** Map showing the region of Brazil where field excursions were made (squared area). Colored circles, and their respective two-letter codes, refer to the Brazilian states where sampling sites are located. **B-D.** Distribution of sampling sites for each bee species within the region shown in A. **E.** Haplotype network used to identify *M. quadrifasciata* subspecies (circle sizes indicate abundance and colors indicate geographic origin).

### 16S ASV metabarcoding

We used in house python scripts to assign the demultiplexed sequencing reads to each bee sample. After primer trimming, 16S rRNA gene sequences were filtered for length (between 1400 and 1600 pb) and quality (maximum expected error = 2). Amplicon Sequence Variants (ASVs) were constructed using DADA2 [33] in R version 4.4.0 with default parameters (except errorEstimationFunction=PacBioErrfun and BAND_SIZE=32). Chimeras were removed from the dereplicated and denoised dataset with the removeBimeraDenovo dada2 function, and then ASVs were assigned to the sequences based on the SILVA database [34] version 138.1 and the dada2 function assignTaxonomy (that uses the RDP Naive Bayesian Classifier algorithm described in Wang et al. [35], with kmer size 8 and 100 bootstrap replicates). The raw sequences of this dataset are available at NCBI under the BioProject <u>PRJNA1307476</u>.

### Phylogenetic analyses

ASVs from the *Melipona* spp. core microbiota were aligned to reference sequences (**Supplementary File 2**) and used for phylogenetic inferences with Geneious Prime 2025.2.1 (Biomatters, Auckland, New Zealand). Alignments were performed using Muscle 5.1 [31] and phylogenies were inferred by maximum likelihood with PhyML [36] using the GTR model with estimated transition/transversion ratios, proportion of invariable sites and gamma distribution parameters.

### Bacterial community analyses

All statistical analyses were performed in R version 4.4.1 [37] using RStudio version 2024.09.1+394 (Posit Software, Boston, Massachusetts), on a dataset from which negative controls and samples with less than 7,000 reads had been removed. Our codes are available at https://github.com/klhaag/metabarcoding-melipona/tree/main/Data_04. To determine whether samples had been sequenced to sufficient depths to capture the true community diversity, rarefaction curves were generated with phyloseq [38] by plotting the number of ASVs observed in each subsample using 7,000 randomizations (“step”=200). Alpha-diversity estimators (Shannon diversity index and Chao1 estimated richness) were also obtained with phyloseq [38]. To test if stingless bees from different species (or subspecies) harbored microbiomes with distinct diversities we used Kruskal–Wallis tests followed by Dunn tests, and correcting for multiple comparisons with the Benjamini-Hochberg method (FDR). For all statistical analyses, values were considered statistically significant when *p*<0.05.

Core microbiomes were inferred based on taxa relative abundance and prevalence, as previously suggested [39]. The top most abundant and prevalent genera were calculated, in order to identify a “common core”. Beta-diversity (Bray–Curtis distance calculated from ASV relative read counts) was used to identify whether the composition of bacterial gut microbiomes differed between species and subspecies. Differences were visualized by non-metric multidimensional scaling (NMDS), and significance was calculated through PERMANOVA using the “adonis2” function from the *vegan* package [40], correcting for multiple comparisons with the FDR method.

Both the host species (or subspecies) and the sampling site were used as factors in the analyses, and omega squared (𝜔^2^) [41] was used to calculate the amount of total variation in microbiota composition explained by each factor (𝜔^2^ was chosen over r^2^ because it is not sensitive to the number of categories within a factor and can thus be compared across factors). For all PERMANOVA tests, analyses were restricted to municipalities where the species, or haplotypes, under comparison were sampled in sympatry, ensuring that observed differences were not confounded by non-overlapping geographic distributions Differential abundance analyses were conducted to identify bacterial genera whose relative representation in *M. quadrifasciata* colonies varied according to (i) mitochondrial haplotype group (sub-species), and (ii) the proximity to *A. mellifera* apiaries. For both comparisons, the filtered 16S rRNA dataset was subset to include only *M. quadrifasciata* samples. Statistical analyses were performed at the genus level using ANCOM-BC2 (Analysis of Compositions of Microbiomes with Bias Correction 2) [42], as implemented in the ancombc2 R package. The models were fitted with “Region” or “HapGroup” as fixed effects, applying a prevalence cutoff of 1% (prv_cut = 0.01), FDR correction for multiple testing, and default bias-correction parameters (pseudo_sens = TRUE, s0_perc = 0.05, struc_zero = TRUE, neg_lb = TRUE). Differentially abundant taxa were defined as those showing significant bias-corrected log-fold changes (|LFC|>0, *p*<0.05). For each taxon, ANCOM-BC2 estimates of log-fold change and standard error were extracted and visualized as bar plots relative to the reference group.

In order to incorporate the interactional aspect of the *M. quadrifasciata* core microbiome, focusing on the correlations of the “common core” with less abundant bacterial taxa, a network analysis was carried out with the package NetCoMi [43]. Data were preprocessed by separating ASVs associated to each host species and grouping them by their genus. Only bacterial genera with more than 5,000 read counts were considered in the analysis. Networks were made by Spearman correlation, as we do not assume linearity. Correlations were tested for statistical significance by bootstrapping 1,000 permutations and adjusted for multiple tests using FDR adjustment. Only statistically significant relationships were considered (*p*<0.05).

### Identification of shared taxa and putative symbiont spillover

To identify bacterial lineages shared among host species, we used ASV-level overlap based on presence/absence matrices of ASVs assigned to each host. The sets of ASVs detected in *A. mellifera*, *M. quadrifasciata*, and *M. mondury* were compared using Venn diagrams. Genera occurring in at least two hosts were defined as shared taxa, while those restricted to one host were considered host-specific. Specifically, 42 genus-assigned ASVs were shared between *A. mellifera* and *M. quadrifasciata*, and among these, 19 were also detected in *M. mondury*. These 42 ASVs were then used for subsequent prevalence and enrichment analyses.

For each ASV and host, prevalence was calculated as the proportion of individual samples positive for that ASV (number of detections divided by total samples per host). Relative abundance was derived from normalized ASV counts per individual, expressed as the fraction of total reads per sample. To prevent zero-inflated distributions and enable median-based comparisons, a small pseudocount (1/12000, approximating one read per library) was added to all abundance values prior to statistical testing. Host bias, defined as the preferential enrichment of an ASV in one host species (the putative donor) relative to another, was evaluated through pairwise differential-abundance comparisons among the three bee species. For each ASV and host pair, we tested prevalence differences using Fisher’s exact tests and abundance differences using Wilcoxon rank-sum tests, with *p*-values adjusted for multiple testing using FDR. An ASV was considered significantly enriched in a host when both prevalence and relative abundance were higher in that host after FDR correction.

## Results

### Full-length 16S rRNA gene amplicon sequencing of the gut microbiota of sympatric *A. mellifera*, *M. quadrifasciata*, and *M. mondury*

We sampled 167 managed colonies of three eusocial bee species (41 *A. mellifera*, 37 *M. mondury* and 89 *M. quadrifasciata*) from a total of 30 locations, covering most of the known geographic range of *M. quadrifasciata* (**Fig. 1**). For three of the 30 locations, all three species were sampled, while in 24 locations, two of the three species were sampled. Of the 89 colonies of *M. quadrifasciata*, 46 and 43 were identified as subspecies *M. q. quadrifasciata* and *M. q. anthidioidis*, respectively. From each colony, one adult worker bee was used for gut microbiota analysis. To obtain high taxonomic resolution beyond the bacterial species level, we used PacBio full-length 16S rRNA gene amplicon sequencing. We obtained on average 14,303 reads per sample and detected a total of 4,376 ASVs with an average of 59 ASVs per sample. A total of 658 ASVs (15%) could not be annotated at the genus level. The largest proportion of unassigned reads at the genus level in our dataset comes from *M. quadrifasciata* (78.4%), followed by *M. mondury* (19.8%) and *A. mellifera* (1.9%). Most of the unassigned reads belong to the Lactobacillaceae (55.5 %, corresponding to 289 ASVs) and Acetobacteraceae (31%, 81 ASVs).

### Host identity and location explain gut microbiota composition across the three sampled bee species

To assess differences in overall community diversity between the three bee species, we first looked at alpha-diversity. Shannon diversity index (SDI) was not significantly different between the three bee species (**Fig. 2A**). However, the Chao1-estimated richness (CER), which emphasizes the richness of rare ASVs, was slightly higher in *M. mondury* as compared to the other two bee species (Kruskal-Wallis X^2^=11.076; *p*=0.0039; **Fig. 2B**). CER also showed a weak correlation with latitude (Spearman’s rho=0.1524; *p*=0.0535), whereas SDI did not (Spearman’s rho=0.0252; *p*=0.7504). No difference in alpha-diversity was observed for the gut microbiota of the two subspecies of *M. quadrifasciata* (SDI, **Fig. 2C**, and Chao, **Fig. 2D**, both *p*>0.05).

**Figure 2.**
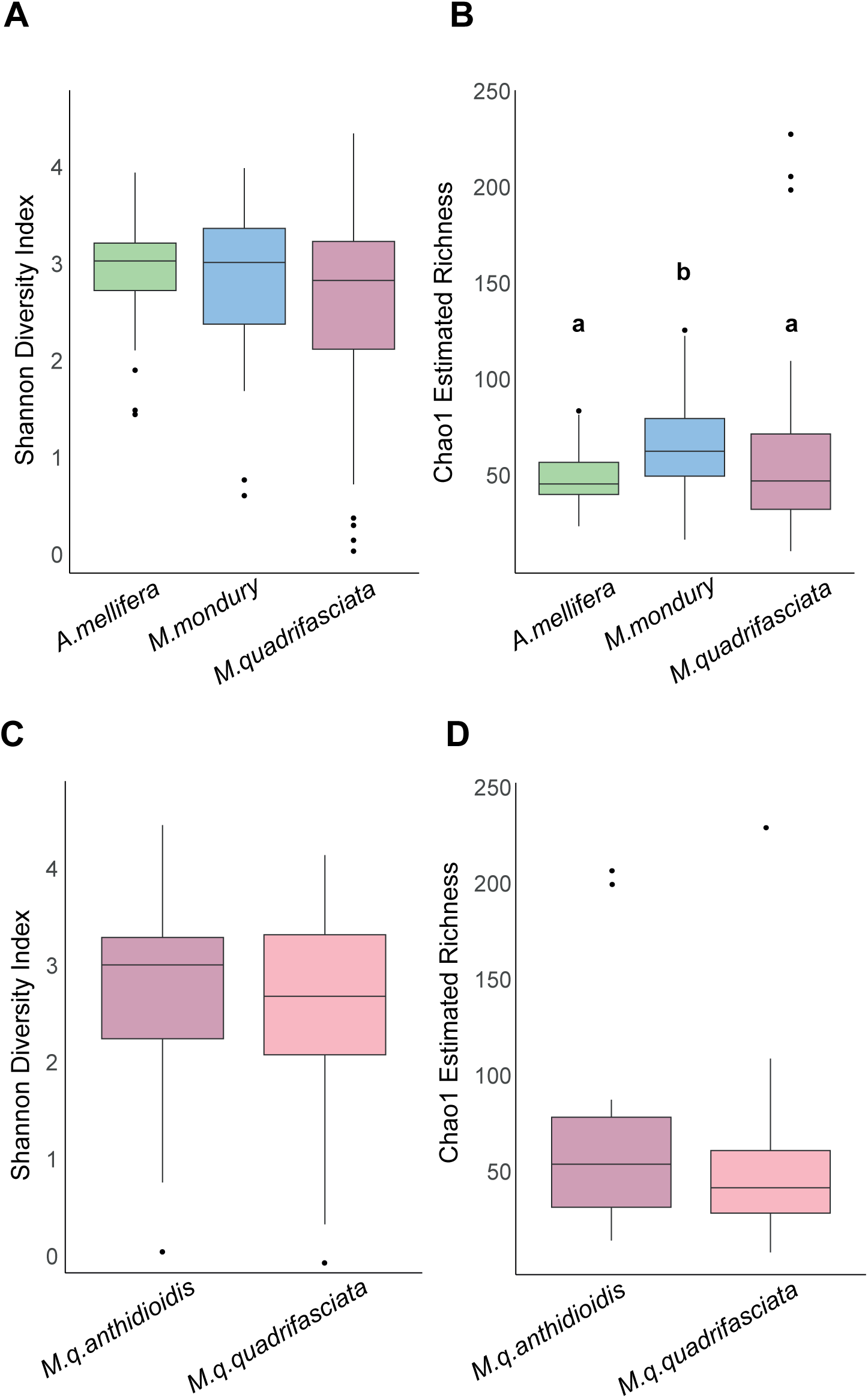
Alpha diversity estimates. Distribution of Shannon diversity index and Chao1 richness estimator obtained for bacterial communities associated to each bee species (**A** and **B**) as well as the subspecies of *M. quadrifasciata* (**C** and **D**).

We next examined patterns of beta-diversity across hosts and habitats. Samples from the three bee species clustered separately in a three-dimensional NMDS ordination of pairwise Bray–Curtis distances (stress=0.1099; **Fig. 3A**). To formally assess the relative contributions of host and environmental factors, we used a two-factor PERMANOVA model including host identity (species or subspecies) and habitat (municipality), as well as their interaction. Importantly, each analysis was restricted to municipalities where all taxa under comparison were sampled, ensuring that observed effects were not confounded by non-overlapping geographic distributions. Across the full dataset, host identity explained approximately 14% of the total variance in community composition (𝜔²=0.1444; *p*=0.001; **Table 1**). This pattern of host specificity remained significant even when either *A. mellifera* or *M. mondury* was excluded, with host identity still explaining ∼9% of the variance in both reduced datasets (𝜔²=0.0953 and 𝜔²=0.0856, respectively; *p*=0.001).

**Figure 3.**
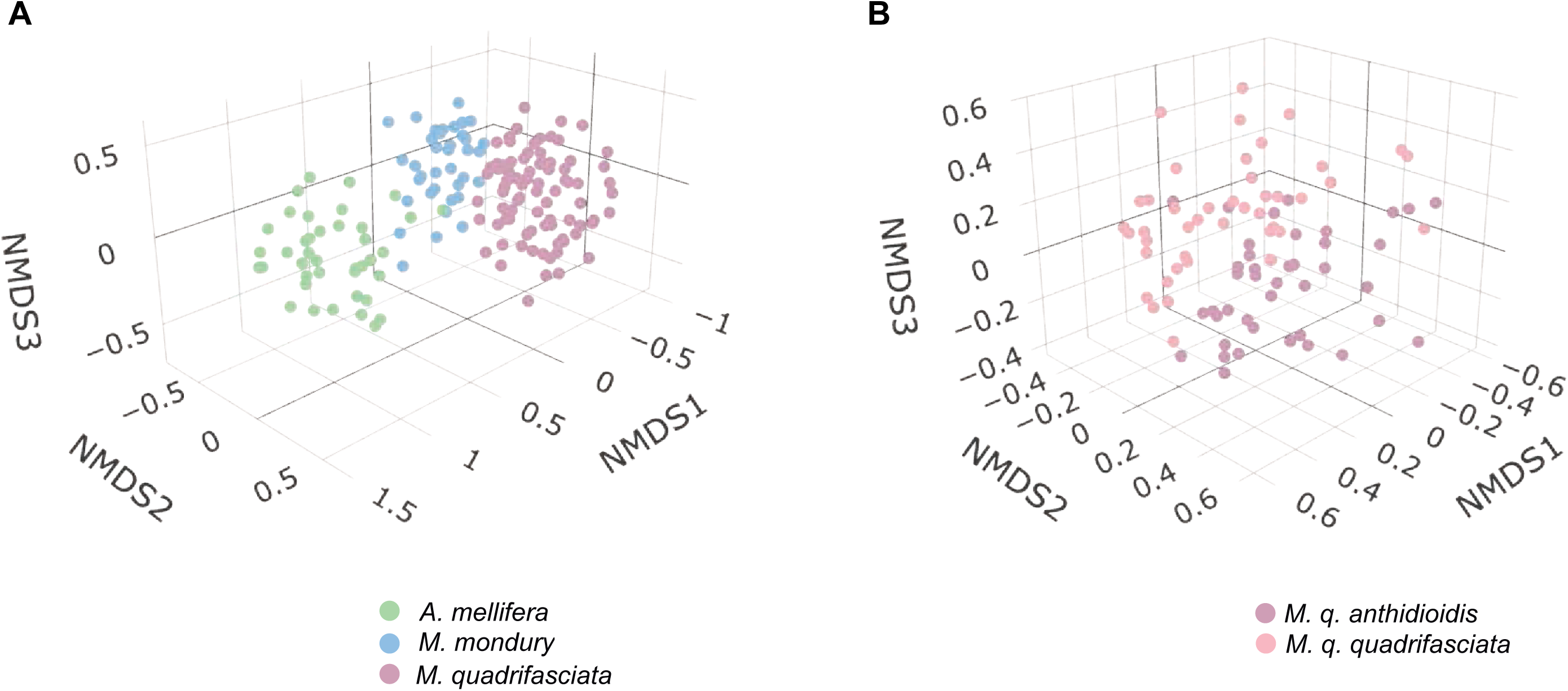
**NMDS.** Three-dimensional Non-metric Multidimensional Scaling analyses of gut bacterial communities. **A.** Analysis ran on the whole dataset, with samples coloured by host species (stress=0.1099). **B.** Analysis ran on *M. quadrifasciata* samples coloured by subspecies (stress=0.1818).

**Table 1.** PERMANOVA of Bray–Curtis distances of gut bacterial communities performed on the whole dataset as well as on different data subsets. The degrees of freedom (DF), sum of squares (SQ), corrected percentage of variation explained by each factor (ω^2^) and the statistical significance (P) are indicated.

| Dataset | | DF | SQ | F | $\omega^2$ | P |
| --- | --- | --- | --- | --- | --- | --- |
| All spp. | Host species | 2 | 2.001 | 2.941 | 0.1444 | 0.001 |
|  | Habitat | 3 | 1.853 | 1.816 | 0.0962 | 0.001 |
|  | Interaction | 3 | 1.748 | 1.713 | 0.0851 | 0.001 |
| *M. quadrifasciata* and *A. mellifera* | Host species | 1 | 3.328 | 8.954 | 0.0856 | 0.001 |
|  | Habitat | 14 | 7.725 | 1.485 | 0.0739 | 0.001 |
|  | Interaction | 14 | 7.638 | 1.468 | 0.0716 | 0.001 |
| *M. mondury* and *A. mellifera* | Host species | 1 | 0.932 | 2.685 | 0.1010 | 0.001 |
|  | Habitat | 3 | 1.828 | 1.755 | 0.1312 | 0.001 |
|  | Interaction | 1 | 0.6311 | 1.818 | 0.0517 | 0.001 |
| *Melipona* spp. | Host species | 1 | 2.867 | 8.482 | 0.0953 | 0.001 |
|  | Habitat | 13 | 7.793 | 1.774 | 0.1241 | 0.001 |
|  | Interaction | 11 | 5.740 | 1.544 | 0.0777 | 0.001 |
| *M. quadrifasciata* | Haplotype | 1 | 1.194 | 3.410 | 0.0279 | 0.001 |
|  | Habitat | 28 | 15.880 | 1.620 | 0.1712 | 0.001 |

Habitat (municipality) also exerted a significant influence on beta-diversity, explaining ∼9.6% of the variance across all species (𝜔²=0.0962; *p*=0.001). The relative contribution of habitat increased when comparisons were restricted to more closely related hosts, reaching ∼12.4% when only *Melipona* species were considered (𝜔²=0.1241; *p*=0.001).

Importantly, the interaction term (host × habitat) was also significant in all analyses (*p*=0.001; **Table 1**), highlighting that the influence of local conditions differs among host taxa. Despite these patterns, there was no evidence that stingless bees microbiota are more environmentally structured than those of honey bees: beta-dispersion analyses revealed no significant difference in the variability of Bray–Curtis distances among hosts (ANOVA; *F*=0.7987; *p*=0.4517). Finally, the gut microbiotas of the two *M. quadrifasciata* subspecies (*M. q. quadrifasciata* and *M. q. anthidioidis*) were distinguishable in NMDS space (stress=0.1818; **Fig. 3B**), with haplotype explaining ∼3% of total variance (𝜔²=0.0279; *p*=0.001).

### Different genera dominate the gut microbiota of *M. quadrifasciata* and *M. mondury*, as compared to *Apis mellifera*

We next identified the core microbiota of each bee species, defining core members as genera present in >70% of individuals and with an average relative abundance of >5% (**Fig. 4**). For the gut microbiota of *A. mellifera,* we identified six core genera given our thresholds: *Lactobacillus*, *Bifidobacterium*, *Gilliamella*, *Snodgrassella*, *Bombilactobacillus*, and *Bartonella*. For the two stingless bee species, *M. mondury* and *M. quadrifasciata*, we identified the same five genera as members of the core microbiota: *Lactobacillus* and *Bifidobacterium* were shared with the core microbiota of *A. mellifera*, while the other three genera, *Apilactobacillus*, *Bombella*, and *Floricoccus*, belonged to the core microbiota of only the *Melipona* hosts. Despite these similarities, the two *Melipona* species differed in the prevalence of the core genera: *Apilactobacillus* was more frequent and significantly more abundant in *M. mondury* (Kruskal–Wallis, *p*=0.0000), whereas *Floricoccus* was more prevalent but not significantly more abundant (*p*=0.7928) in *M. quadrifasciata* as compared to *M. mondury*.

**Figure 4.**
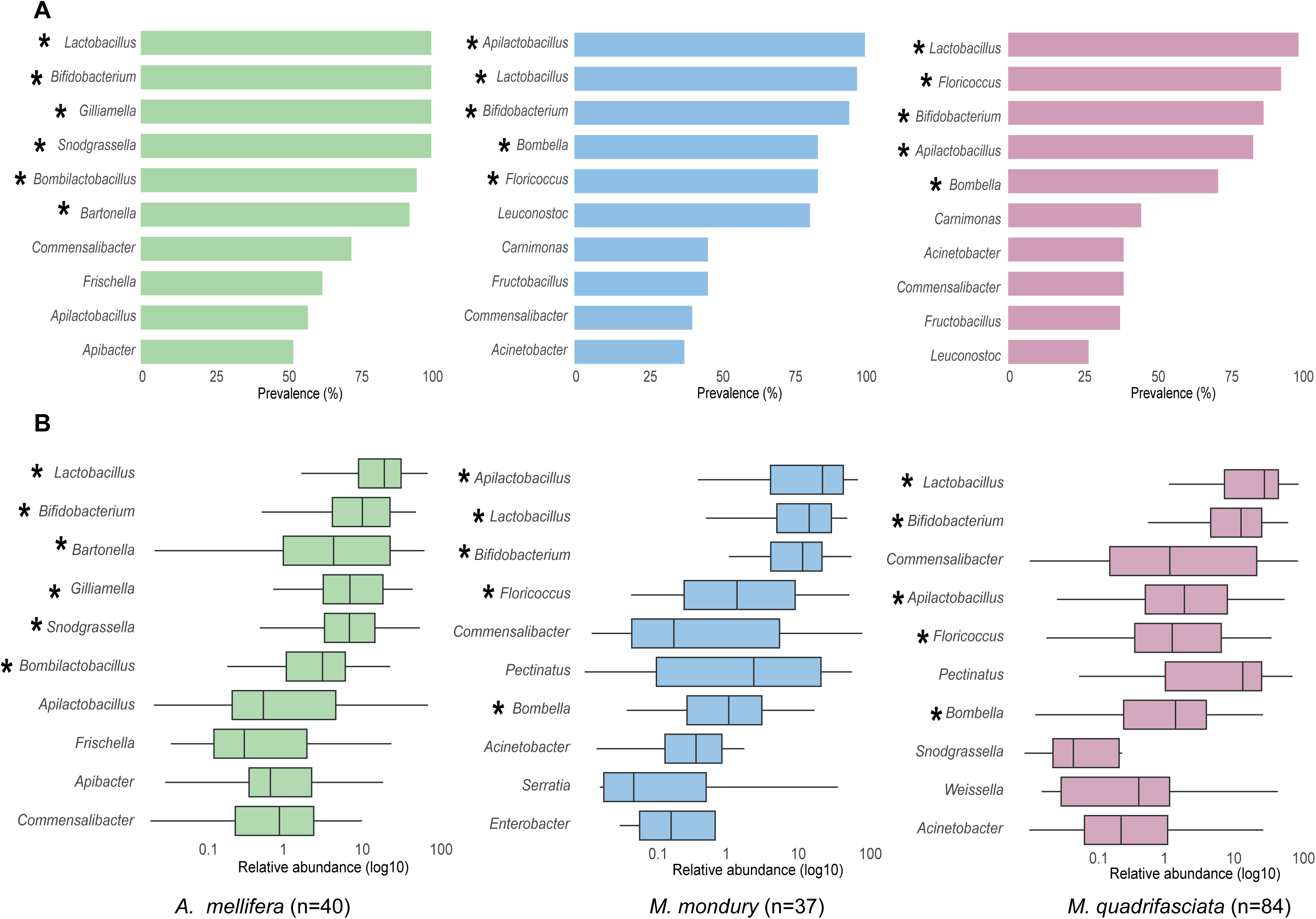
The core genera of the gut microbiota across the three bee species. The ten bacterial genera with the highest **(A)** prevalence and **(B)** relative abundance estimates are shown in descending order of their mean values. Asterisks indicate the core microbiota members of each bee species according to our criteria (see text for details).

### Differences in non-core members explains variation in community composition within *M. quadrifasciata*

Although genetic differentiation within *M. quadrifasciata* only marginally explained Bray–Curtis gut community distances (**Table 1**), we observed clear differences in microbiota composition between the two subspecies of *M. quadrifasciata* in the NMDS analysis. To explore these differences, we performed a differential abundance analyses, and found that the gut microbiota of *M. q. quadrifasciata* was enriched in non-core microbiota members, specifically *Weissella* (β=3.7454, SE=0.2233, *p*=0.0000), *Snodgrassella* (β=3.2946, SE=0.2526, *p*=0.0000), and *Commensalibacter* (β=1.7477, SE=0.4459, *p*=0.0021) compared with *M. q. anthidioidis* (**Fig. 5A**). Because *Snodgrassella* has been reported to be mostly absent from the genus *Melipona* [24], and to test whether habitat could influence its presence via spillover from nearby honey bees, we repeated the analysis comparing colonies kept with versus without *A. mellifera*. Sites where *M. quadrifasciata* co-occurred with honey bees showed enrichment of *Snodgrassella* ASVs, though less pronounced than the subspecies-level contrast (**Fig. 5B**). Since *A. mellifera* colonies co-occurred with 64% of *M. q. quadrifasciata* but only 37% of *M. q. anthidioidis* samples, part of the apparent effect of *A. mellifera* presence on bacterial abundance may be attributable to host genotype.

**Figure 5.**
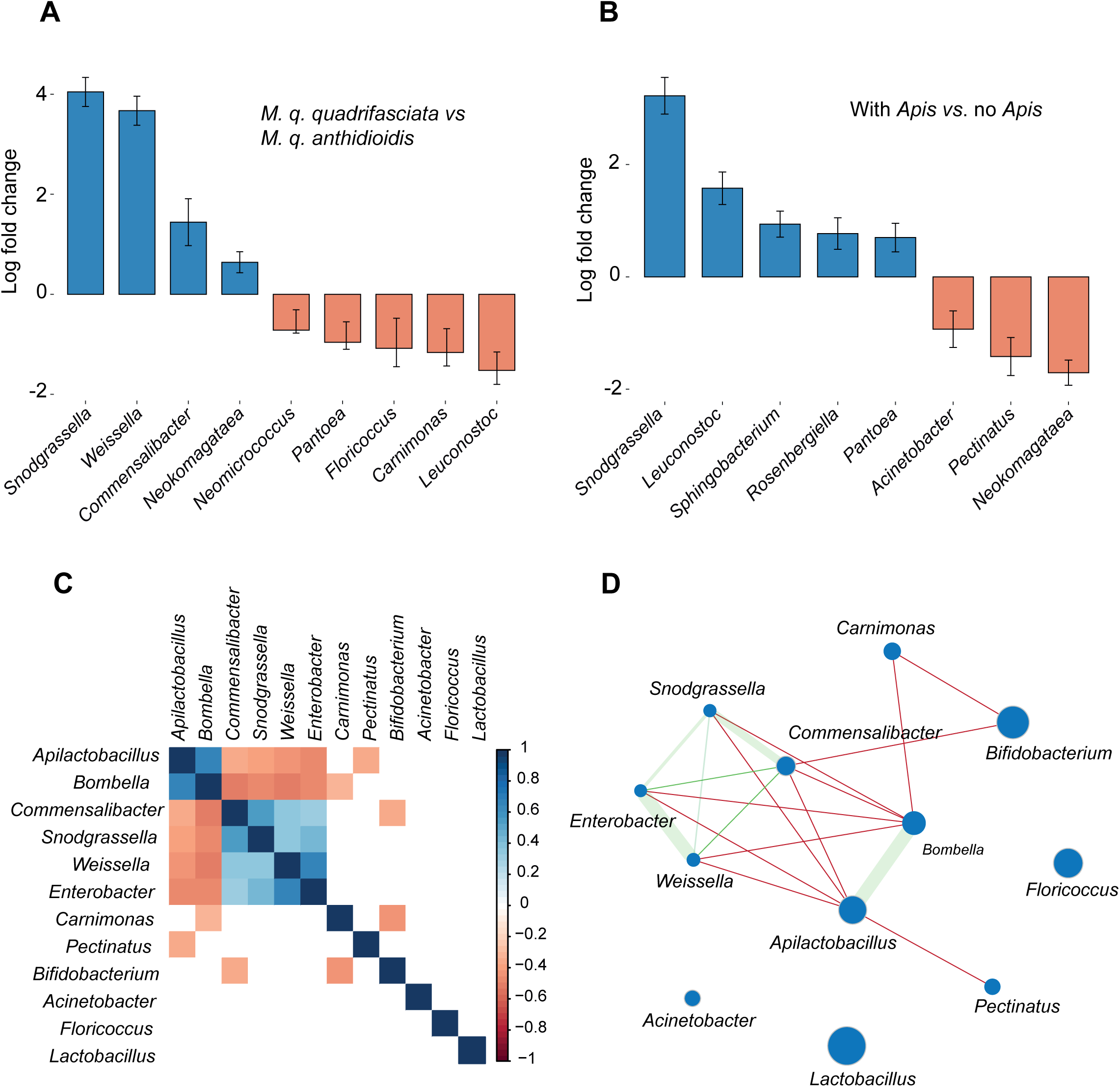
Analyses of bacterial relative abundance in the *M. quadrifasciata* microbiome. **A.** Differential abundance of bacteria between subspecies. Logfold change indicates the amount of enrichment in the number of reads from a genus in *M. q. quadrifasciata* relative to *M. q. anthidioidis*. **B.** Differential abundance of bacteria between sites with and without *Apis*. Logfold change indicates the amount of enrichment in the number of reads from a genus in sites where *M. quadrifasciata* is kept with *A. mellifera* relative to sites without *Apis*. **C.** Matrix of Spearman correlations between relative abundances of bacteria with at least 5,000 reads in the *M. quadrifasciata* dataset. **D.** Network inferred from the statistically significant Spearman correlations. Red and green lines indicate negative and positive correlations, respectively. The width of the green line indicates the intensity of positive correlation.

To further explore the patterns of relative abundance among non-core taxa in *M. quadrifasciata*, we constructed a co-occurrence network that revealed several significant associations (Spearman’s correlation; *p*<0.05; **Fig. 5C–D**). Core bacteria displayed strong positive correlations, particularly between *Apilactobacillus* and *Bombella* (ρ=0.66), and between *Snodgrassella* and *Commensalibacter* (ρ=0.54). A positive association was also observed between *Weissella* and *Enterobacter* (ρ=0.65). In contrast, *Apilactobacillus* and *Bombella* showed consistent negative correlations with *Snodgrassella*, *Commensalibacter*, *Weissella*, and *Enterobacter* (ρ ranging from – 0.35 to –0.52), while *Bifidobacterium* correlated negatively with *Commensalibacter* and *Carnimonas* (ρ=–0.35 and –0.44, respectively). Overall, these patterns indicate that core genera tend to co-occur with each other but are negatively associated with non-core genera sporadically present in the gut microbiota of *M. quadrifasciata*.

### Evidence for gut bacteria transfer between the three analyzed bee species

Because all three bee species were sampled from the same geographic regions, we investigated the potential for gut symbiont transfer among them. Although community-level analysis indicated that the overall gut microbiota composition is host specific (**Fig. 3A**), individual symbionts may occasionally be transferred across species boundaries. The fact that honey bee–specific core members were more prevalent in *M. quadrifasciata* individuals maintained in close proximity to honey bee colonies than in others supports this idea (**Fig. 5B**).

In line with this, we found that 238 of 4,275 ASVs are shared among the three species: 19 were present across all three, 167 were shared between the two *Melipona* species, and 52 occurred between *A. mellifera* and one *Melipona* species (**Fig. 6**). The most frequently shared ASVs belonged to *Apilactobacillus* and *Bombella* and were predominantly shared between the *Melipona* species. ASVs shared with *A. mellifera* belonged to various genera, including five of the six honey bee core members (*Lactobacillus*, *Bartonella*, *Bombilactobacillus*, *Gilliamella*, *Snodgrassella*).

**Figure 6.**
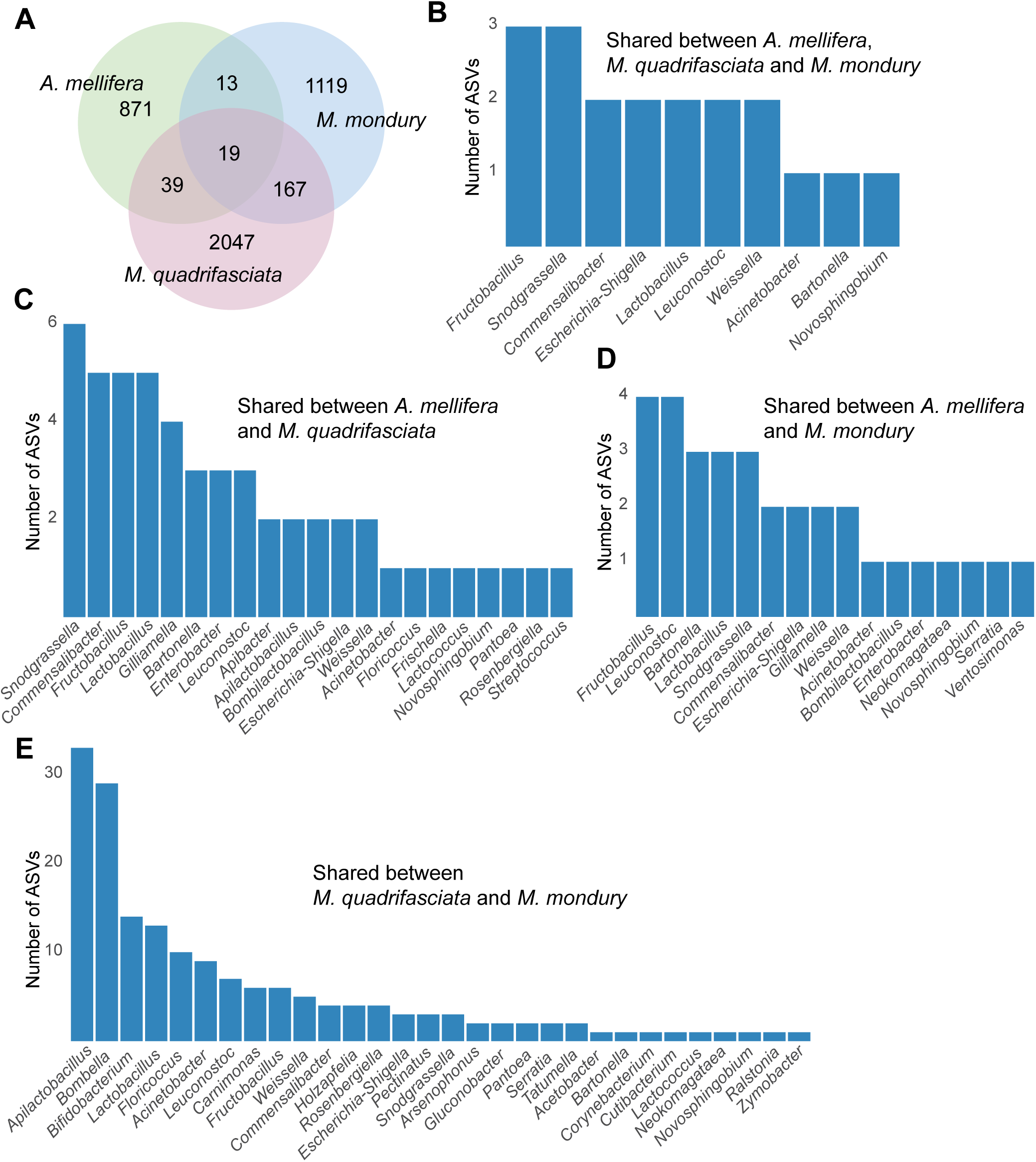
Shared ASVs. **A.** Venn diagram indicating the number of exclusive and shared ASVs per host species. **B-D.** Distribution of the number of shared ASVs per bacterial genera.

To investigate how the shared taxa were distributed across the different bee hosts, we focused on 42 genus-assigned ASVs that were detected in both *A. mellifera* and *M. quadrifasciata* (**Fig. 7A–C, Supplementary File 3**). Of these, 19 were also detected in *M. mondury*, allowing us to compare not only between honey bees and stingless bees, but also between two closely related stingless bee species. Most ASVs occurred at low prevalence in all hosts, but a subset showed clear and statistically supported host preference. For example, ASV2 and ASV40, assigned to *Apilactobacillus/Lactobacillus*-like lineages, were present in 50% and 57% of *M. quadrifasciata* individuals, respectively, but were rarely detected in *A. mellifera* (≤7.5%; Fisher’s exact test, *p*<0.05). These ASVs also reached higher relative abundance in *M. quadrifasciata* than in honey bees (Wilcoxon test, *p*<0.05). ASV36 followed the same pattern: it occurred in ∼39% of *M. quadrifasciata* workers but was rarely present in *A. mellifera* samples (2.5 %), and was not recovered from *M. mondury* at comparable frequency (Fisher’s exact test, *p*<0.05). These patterns indicate that some of the shared ASVs are predominantly found in samples of *M. quadrifasciata* and only sporadically occur in honey bees.

**Figure 7.**
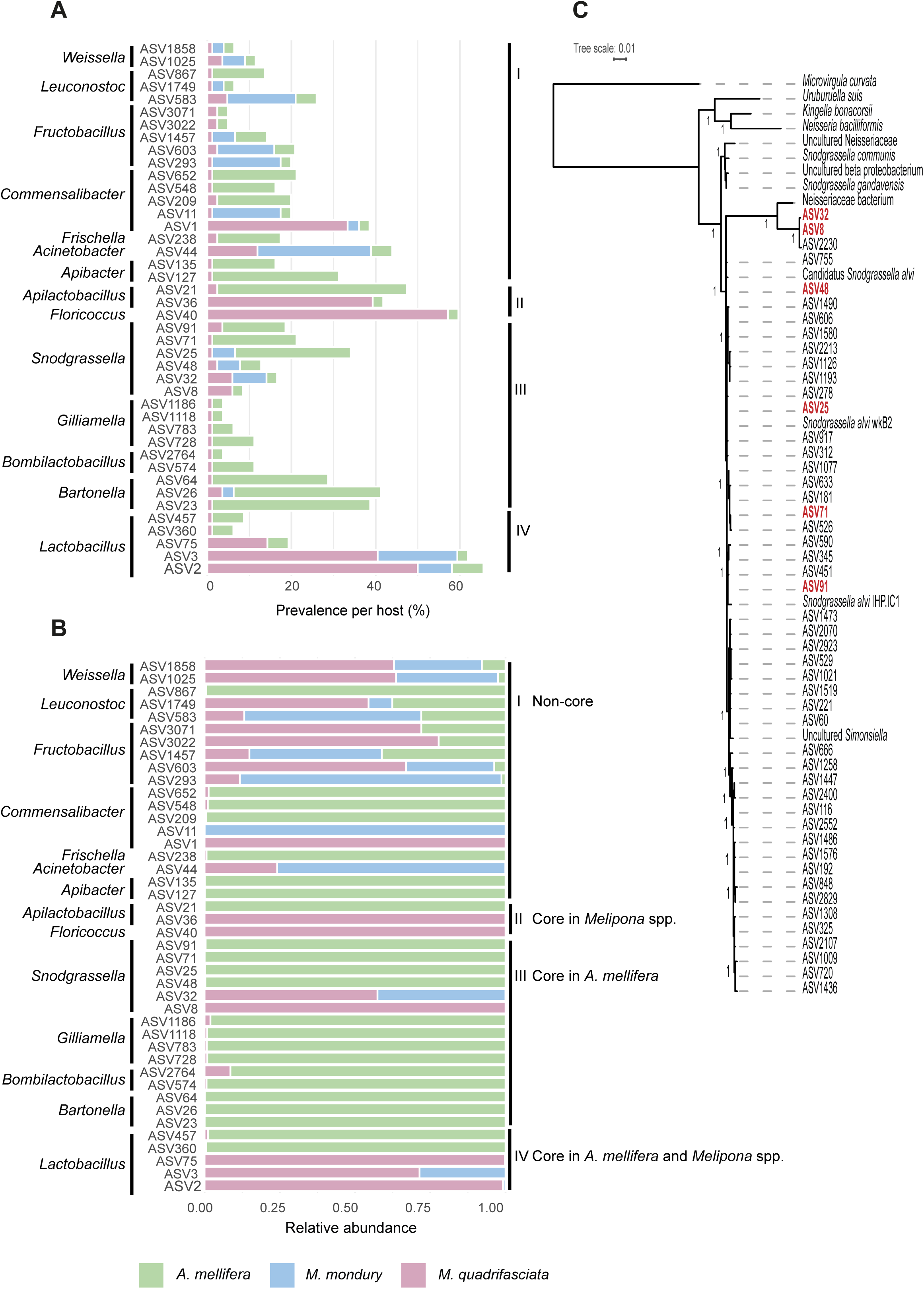
Donor host species of shared ASVs. **A–B.** Relative prevalence and relative abundance of ASVs shared between *A. mellifera* and *M. quadrifasciata*. For each host species, relative prevalence was calculated as the proportion of samples in which a given ASV was detected, normalized by the total number of samples for that host. Relative abundance was calculated as the proportion of total reads assigned to an ASV in each host, relative to the total number of reads from that ASV across the three host species. **C.** Phylogeny of *Snodgrassella* ASVs. ASVs shared between *M. quadrifasciata* and *A. mellifera* are shown in red; the triangle highlights a divergent clade containing *Melipona*-specific ASVs. Shimodaira-Hasegawa-like branch support values > 0 are shown.

However, the inverse pattern was also observed. In particular, ASVs assigned to canonical honey bee symbionts such as *Snodgrassella, Gilliamella/Commensalibacter*, and *Bartonella* were enriched in *A. mellifera* as compared to *M. quadrifasciata* samples. For instance, ASV21 was detected in 45% of *A. mellifera* individuals but only 2.4% of *M. quadrifasciata*; ASV23 in 37.5% *vs* 1.2%; ASV127 in 30% *vs* 1.2% (Fisher’s exact test, all *p*<0.05). The same ASVs also reached higher within-host abundance in *A. mellifera* than in *M. quadrifasciata* (Wilcoxon test, *p*<0.05). Many of these honey bee– biased ASVs were either absent or extremely rare in *M. mondury*, suggesting that these lineages are not broadly established across *Melipona*, but only sporadically make it into *A. mellifera*.

Including *M. mondury* in the comparison further showed that not all *Melipona* hosts carry the same set of ASVs. Among the 19 ASVs that were found in *M. mondury*, a few low-prevalence taxa (for example *Commensalibacter* ASV11 and *Fructobacillus* ASV293) were detected in ∼16% of *M. mondury* samples but in only ∼1% of *M. quadrifasciata*, and they reached higher relative abundance in *M. mondury* (Fisher’s exact test and Wilcoxon test, *p*<0.05). By contrast, high-prevalence *M. quadrifasciata* ASVs such as *Lactobacillus* ASV2 and *Floricoccus* ASV40 were either absent or detected only sporadically in *M. mondury*. This indicates that even between these two closely related *Melipona* species, shared gut bacteria usually have a predominant host species.

### A divergent ASV of *Snodgrasella* shared between stingless bees and honeybees

We examined the shared ASVs of *Snodgrassella* in more detail, because this genus is considered a core symbiont of honey bees but has been proposed to be mostly absent in neotropical stingless bees, in particular the genus *Melipona* [17, 24]. Some of the shared *Snodgrassella* ASVs (*e.g.* ASV25, ASV71) were significantly more prevalent in *A. mellifera* than in *M. quadrifasciata* (up to 20% *vs* ≤3.6% of individuals; Fisher’s exact test, *p*<0.05) and/or showed higher abundance in *A. mellifera* (ASV25, ASV71. ASV91; Wilcoxon test, *p*<0.05). They also were nearly identical to the 16S rRNA gene sequence of *S.alvi* wkB2 (Figure 7), the type strain of this *Snodgrassella* species widely distributed in *A. mellifera* [44]. Interestingly, two additional *Snodgrassella* ASVs shared between *A. mellifera* and *M. quadrifasciata* (ASV8, ASV32) were clearly distinct, sharing only 96% sequence identity in the 16S rRNA gene with those of *S. alvi* and forming a divergent deep-branching clade (**Fig. 7C**). Despite their low prevalence and lack of significant enrichment in either stingless bee host relative to the honey bee samples, these ASVs still reached remarkably high abundances in stingless bee samples from three different locations (see **Supplementary File 4**). Moreover, a *Snodgrassella-*like symbiont previously isolated from the stingless bee species *Frieseomelitta varia* [17] clustered together with ASV8 and ASV32 (**Fig. 7C**) suggesting that this clade is specific to stingless bees.

## Discussion

Previous studies have shown that stingless bees of the genus *Melipona* harbor specialized gut microbial communities that resemble those of other social bees but remain clearly distinct from the well-characterized microbiota of the honey bee, *Apis mellifera* [17, 19, 20, 24, 45, 46]. However, the extent to which the gut microbiota of *Melipona* varies across closely related species, sub-species, or individual bees, especially in comparison to that of honey bees, remains unknown. Additionally, it is unclear whether microbial spillover can occur between stingless bee species and honey bees that are kept in close proximity. Here, we addressed these questions by sampling individual adult bees of *M. quadrifasciata*, *M. modury*, and *A. mellifera* from a total of 167 colonies across the geographic range of *M. quadrifasciata* in Brazil. By sampling all three bee species in the same region, we were able to compare their gut microbial diversity while controlling for environment.

In summary, our results show that both *Melipona* species possess host-specific gut microbial communities that are taxonomically distinct from those of honey bees, but exhibit similar levels of within-and between-host diversity. We find evidence for spillover of core microbial members between the three bee species; however, these typically represent minor components of the gut communities.

In contrast to previous studies that characterized the gut microbiota of stingless bees using partial 16S rRNA gene amplicon sequencing of the V3-V4 region, we employed PacBio full-length 16S rRNA gene sequencing. This approach provides higher taxonomic resolution of the resulting ASVs, which enabled us to distinguish closely related bacteria and assess differences in community composition below the bacterial species-level within and across the analyzed bee species.

Our study confirms previous findings that the gut microbiota of social bees is host-specific [15, 17]. *M. quadrifasciata*, *M. mondury*, and *A. mellifera* each harbored distinct gut communities (see also **Supplementary File 5**). Notably, we detected differences between the two subspecies of *M. quadrifasciata* as well, despite both harboring the same bacterial genera, highlighting the high taxonomic resolution provided by full-length 16S rRNA gene amplicon sequencing. However, our PERMANOVA analyses also revealed that habitat significantly contributes to gut microbiota structure. In fact, habitat explained more of the observed variance than host identity when comparing the microbial communities within *Melipona* samples, particularly between the two subspecies of *M. quadrifasciata*. These results demonstrate that, while host specificity remains a major determinant of gut microbiome composition, environmental context is a key source of variation among closely related hosts. Yet, our analysis provides no evidence that environment plays a more important role in structuring the gut microbiota of stingless bees as compared to honey bees: *Melipona* spp. showed neither higher alpha-diversity (Shannon diversity index and Chao1 estimated richness) nor greater beta-diversity dispersion (Bray–Curtis distances) than honeybees across the sampled sites. Only *M. mondury* exhibited slightly higher Chao1 estimates, but the absence of samples from its southernmost distribution and the weak correlation between richness and latitude may have biased comparisons. Previous studies have documented associations between gut bacterial diversity and latitude in both honey bees and stingless bees [47, 48].

Our broad sampling enabled us to identify the “core” genera dominating the gut microbiota of the two stingless bee species. We applied an arbitrary cutoff to define core taxa. Therefore, future studies, *e.g.* when conducted across seasons, may come to different results. Our analysis nonetheless offers a quantitative assessment of the most prevalent bacterial groups within the microbiota of the two stingless bee species as compared to honey bees sampled in the same region. The two *Melipona* species shared the same five core genera; however, only two of these (*Lactobacillus* and *Bifidobacterium*) were also part of the core microbiota of honey bees. Some of the other core genera identified in *Melipona*, such as *Bombella* and *Apilactobacillus*, are also repeatedly detected in honey bees but appear to be less dominant in those hosts.

Conversely, Proteobacterial core members typical of honey bees, particularly *Gilliamella* and *Snodgrassella*, are rare in *Melipona* species, although not entirely absent, confirming previous findings [17]. Overall, these results indicate that the gut microbiota of *Melipona* is dominated by a distinct set of bacterial core genera compared to honey bees, supporting the idea that the gut microbiota of social bees has undergone substantial compositional changes during evolution.

Despite the host-specific nature of the gut microbiota at the community level in the three bee species, we detected a non-negligible proportion of ASVs (6% of all ASVs) that were present in samples from different host species. The larger fraction of these ASVs was shared between the two *Melipona* species than between *Melipona* and honey bees, suggesting that host phylogeny and ecology may act as barriers to host switching. Although we cannot exclude the possibility that bacteria with the same ASV differ in other parts of their genomes, and hence are evolutionary divergent, these findings suggest that some gut bacteria have been recently transferred between bee species [37, 38]. We can also not rule out the chance that some of the detected shared ASVs are artifacts of amplicon sequencing or result from cross-contamination during sample processing. However, the relatively high abundance of several shared ASVs across multiple samples makes such explanations unlikely.

Shared ASVs may represent non-specific, environmental bacteria that are occasionally acquired by different social bee species (*e.g.,* during flower visits), or they could be specialized gut symbionts that are common to multiple host species or primarily associated with one species but occasionally spill over into others. Our enrichment analysis of the 42 genus-assigned ASVs shared between *A. mellifera* and *M. quadrifasciata* showed that many of them belonged to core genera and that they were typically much more prominent in one host versus another. A particularly interesting case was that of *Snodgrassella*. This genus is typically rare in *Melipona* stingless bees [17, 24]. However, we detected several *Snodgrassella* ASVs in our *Melipona* samples that were shared with honey bees. Some of these ASVs clustered within a honey bee– specific clade of *Snodgrassella*; and although their abundances were generally low, their presence indicates that this core gut symbiont can spill over from introduced honey bees into native stingless bees. In other cases, however, the shared ASVs belonged to a divergent clade of *Snodgrassella* specific to stingless bees, suggesting that bacterial exchange can occur in both directions. Overall, these results suggest that occasional spillover of core members from their native host into a non-native host does occur. Although the generally low prevalence and abundance of these ASVs suggest that such bacteria often fail to establish stable associations in the secondary host, likely due to competition with the resident microbiota, our results demonstrate that opportunities for host switching are present. Human beekeeping practices such as mixed apiaries, colony manipulations, or supplemental feeding of stingless bee colonies with honeybee products are likely to increase such opportunities. That stingless bees are susceptible to such environmental factors has been shown in the case of Australian stingless bees, which exhibited changes in gut microbiota composition after their colonies were translocated for pollination services [49, 50]. Moreover, recent studies have shown that the gut microbiota of social bees have not strictly co-diversified with their host, but that symbiont gain, loss, and exchanges have occurred. This suggests that such ecological opportunities of gut microbiota exchange can result in novel, evolutionary stable associations.

## Conclusion

Our study provides a comprehensive strain-level view of the gut microbiota of two important stingless bee species in Brazil and shows that, although *Melipona* communities are compositionally distinct, human-mediated beekeeping creates ecological contact zones that allow gut symbionts to cross species boundaries. We found no evidence that stingless bee microbiota are inherently more environmentally assembled than those of honey bees, but we did detect widespread ASV sharing, including all *A. mellifera* core taxa except *Bifidobacterium,* and discovered divergent *Snodgrassella* lineages thriving in some stingless bee colonies. Our results suggest that management practices such as artificial feeding, colony transport, and mixed apiaries may promote spillover. The ability of such occasionally transferred bacteria to establish themselves in non-native stingless bee hosts and to influence host–microbe or microbe–microbe interactions in the gut warrants further investigation. Whole-genome sequencing and functional studies will be essential to further elucidate the causes and consequences of gut microbiota variation in stingless bees of the genus *Melipona*, and specifically to understand how anthropogenic influences reshape bee gut communities and affect pollinator health.

## Acknowledgements

This work was funded by the Swiss National Science Foundation through a Spirit grant (grant number 189496 to P.E.), an SNSF Consolidator grant (’GLOBEE,’ grant number 213860 to P.E.), a scientific exchange grant (IZSEZ0_224300 to P.E. and K.L.H.) and by the Fundação de Amparo a Pesquisa do Estado do Rio Grande do Sul through a PQG Grant (19/2551-0001860-6 to K.L.H). We thank to the following beekeepers for providing us the bees analyzed in our study: Alexandre Nunes (Meliponário Sabor e Saúde, RJ), Alexandre Piccini (EMATER, RS), André da Silva Xavier (UFES, ES), André Marildo Sezerino, Elson Juarez da Silva (INCAPER, ES), Evald Gossler, Fábio Junior V. Calegari, Glaucio Canpanatti (Amesampa, SP), Gledison J. T. de Souza, Hércules Birchner (Jardim de Mel, ES), João Robaski Meregali, Jorge Luis Palu, Leonardo Pereira (Meliponário LP, ES), Manoel A. de Souza, Marcelo da Silva Campos (Recanto das Nativas, RJ), Marcelo Passos Gonçalves (Produtos da Colmeia), Marina Souza Cunha (UFRRJ, RJ), Miguelângelo Z. Arboitte, Paulo Rocha (Apiários e Meliponários Panabee, SP), Raul Maciel Gulart, Simone Prado (EMBRAPA/SP), Vinicius Deloel Monteiro Gomes, Viriato Dutra Neto (Meliponário Farroupilha, RS). We thank the Lausanne Genomics Technology Facility (GTF) team at the University of Lausanne for performing the high-throughput sequencing for the experiments.

## SUPPLEMENTARY DATA

**Supplementary File 1.**
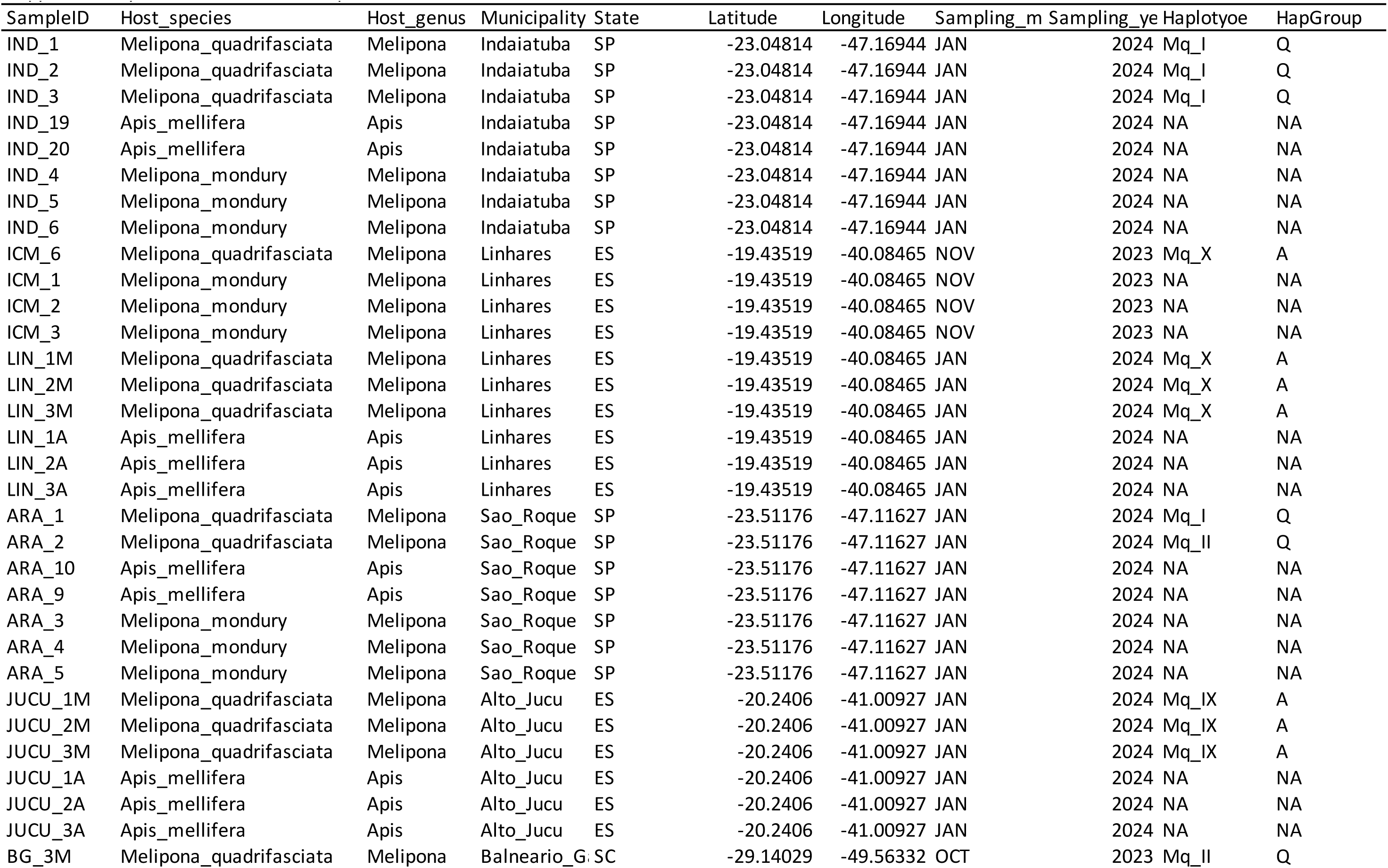
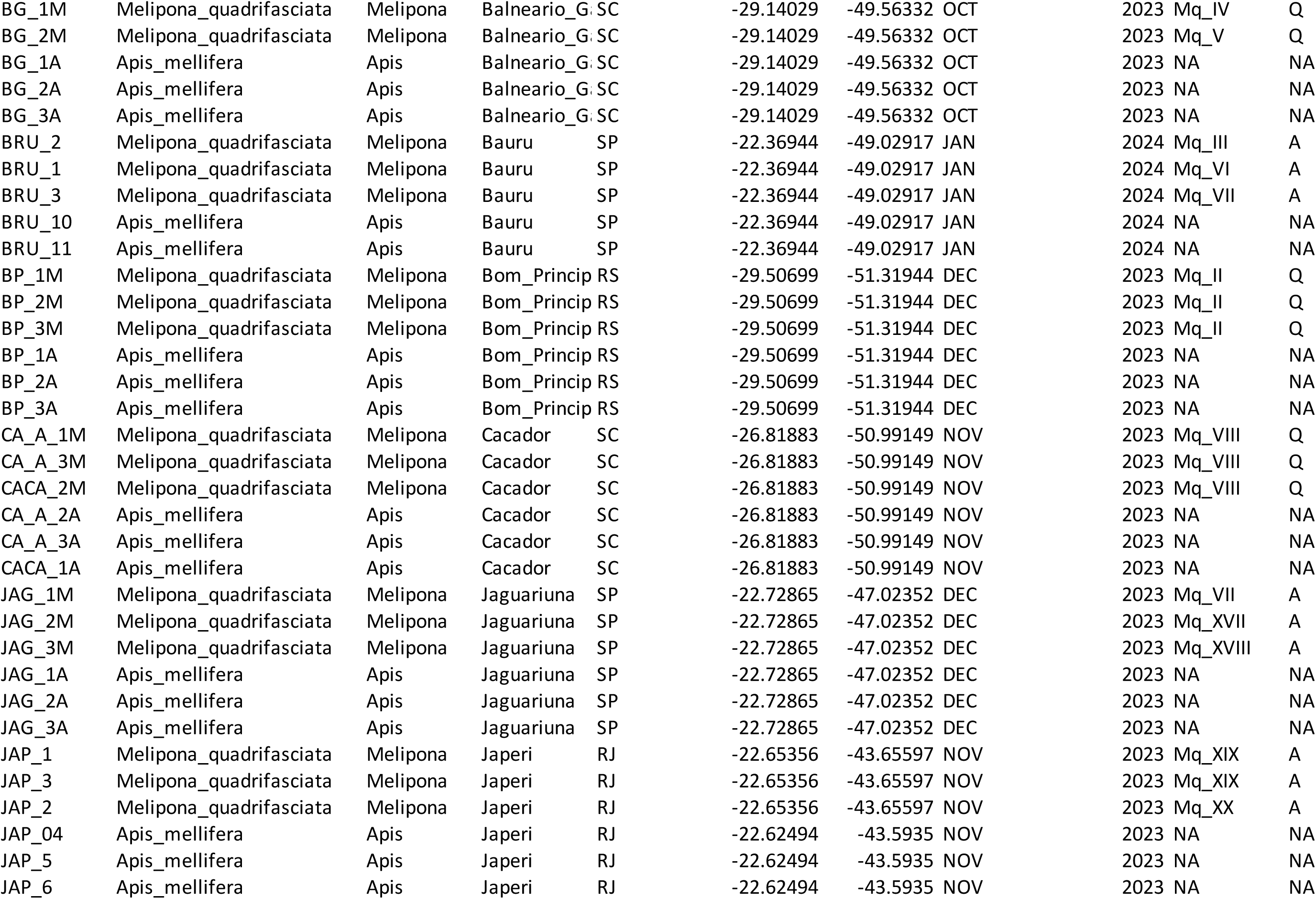
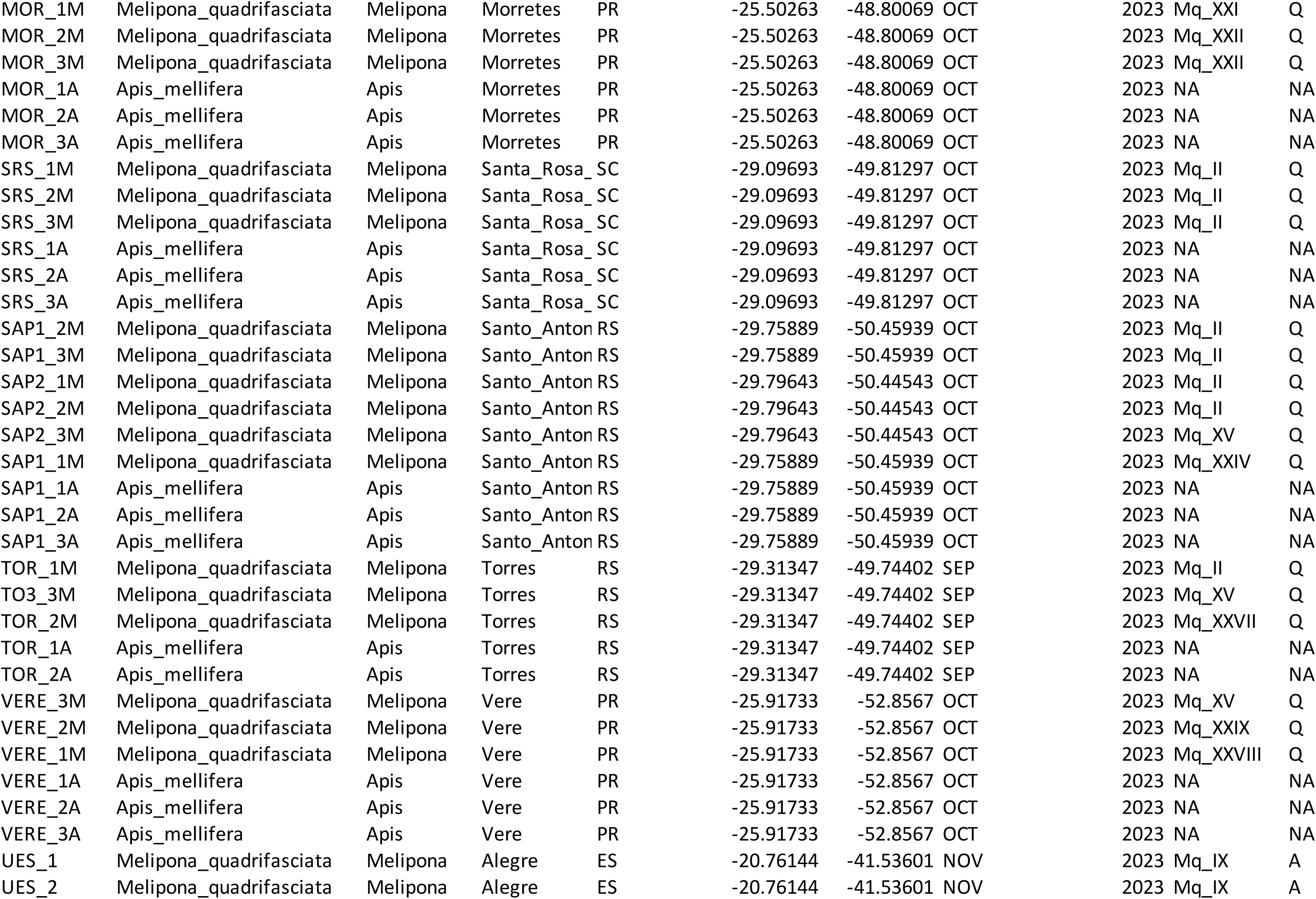
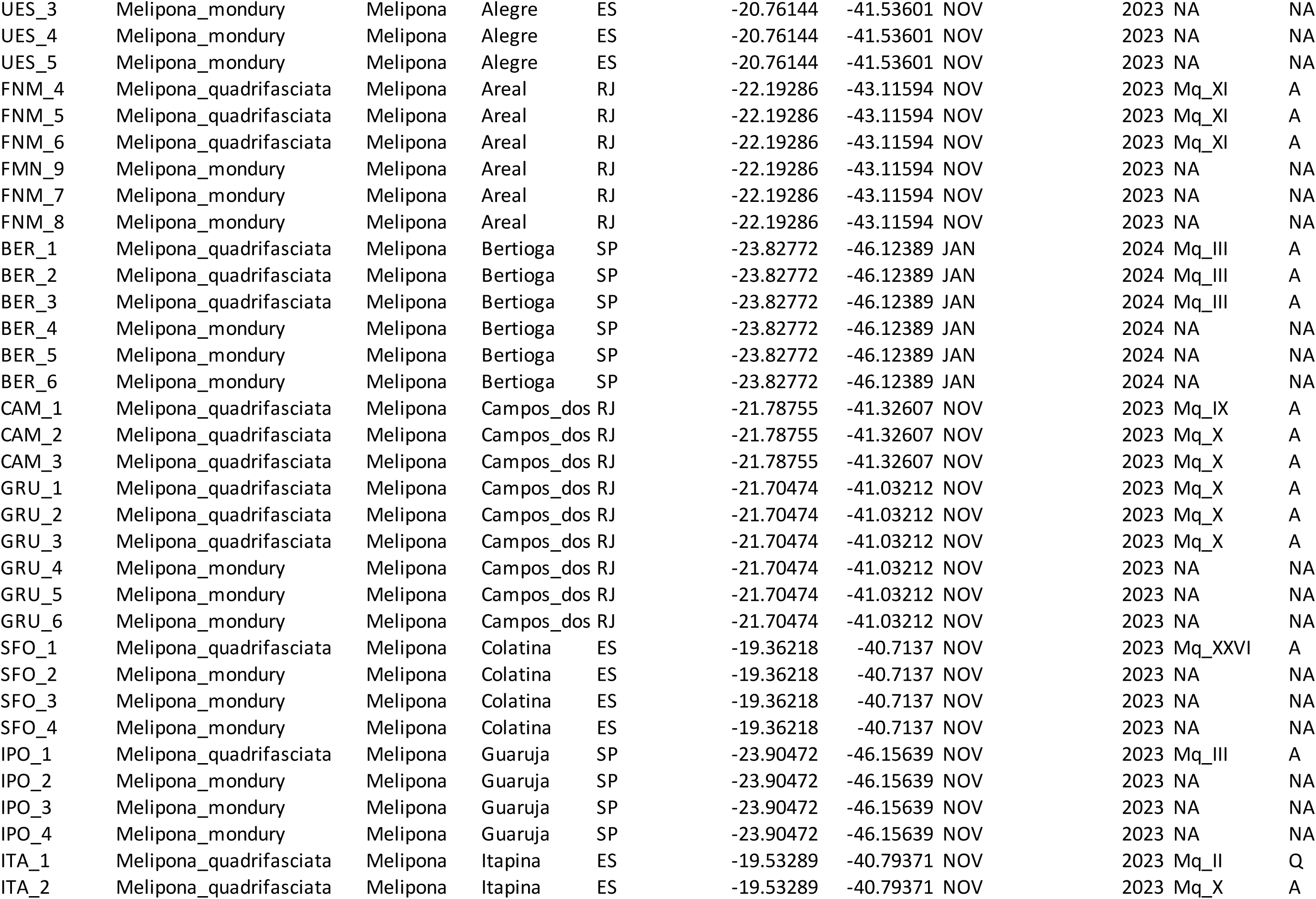
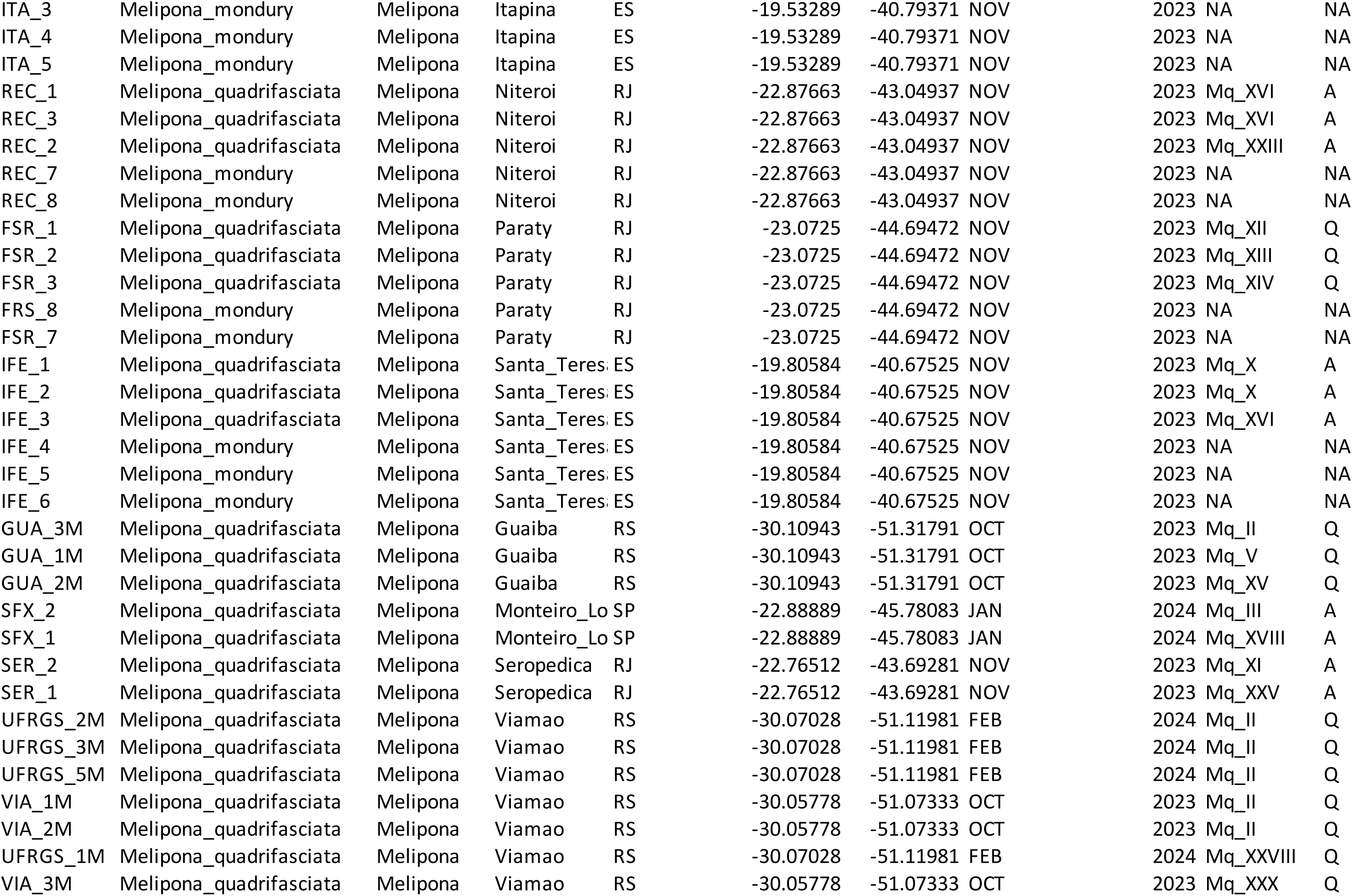
List of bee samples and their associated metadata.

**Supplementary File 2.**
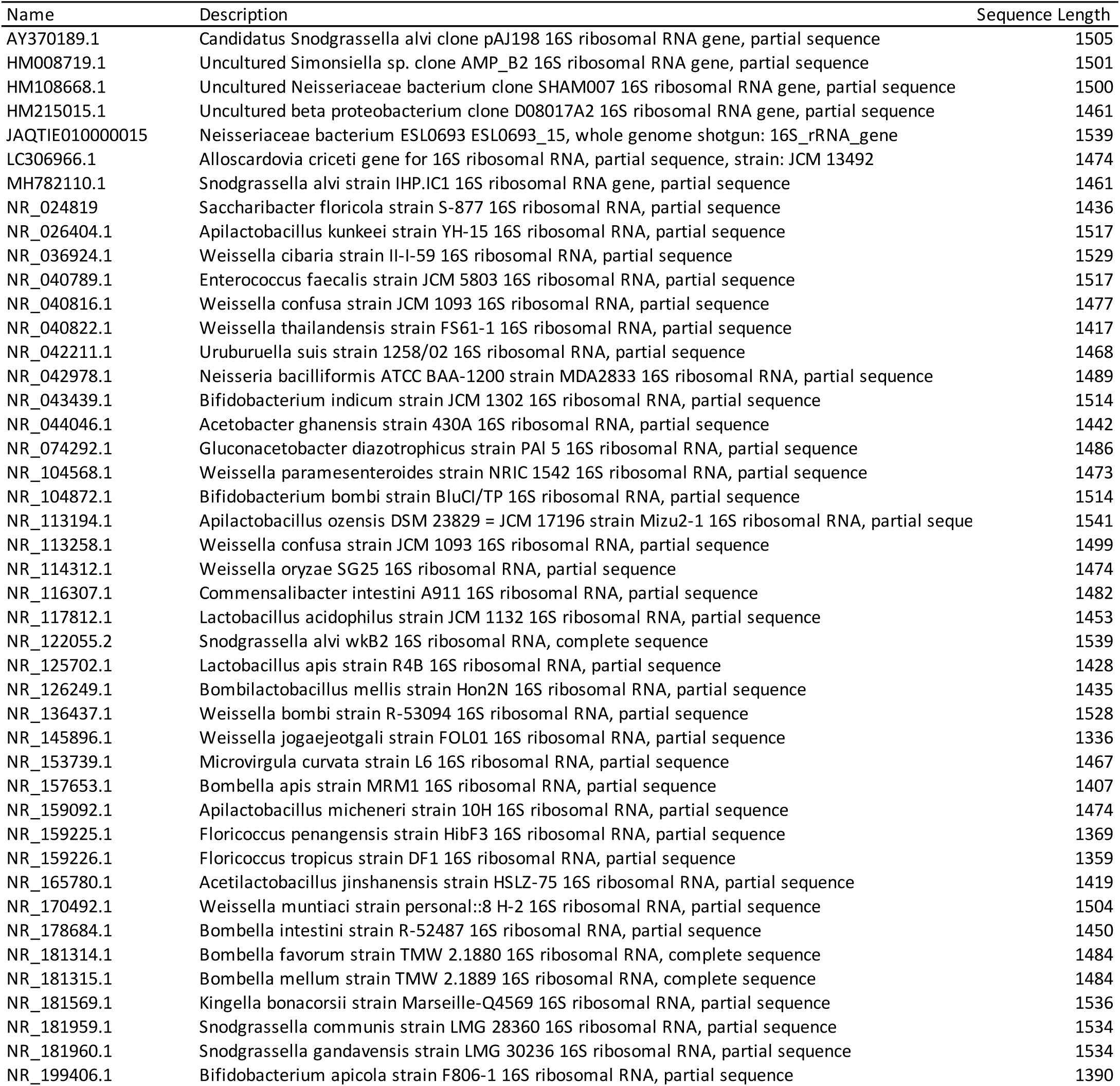
GenBank accession numbers of the reference sequences used in the phylogenetic analyses.

**Supplementary File 3.**
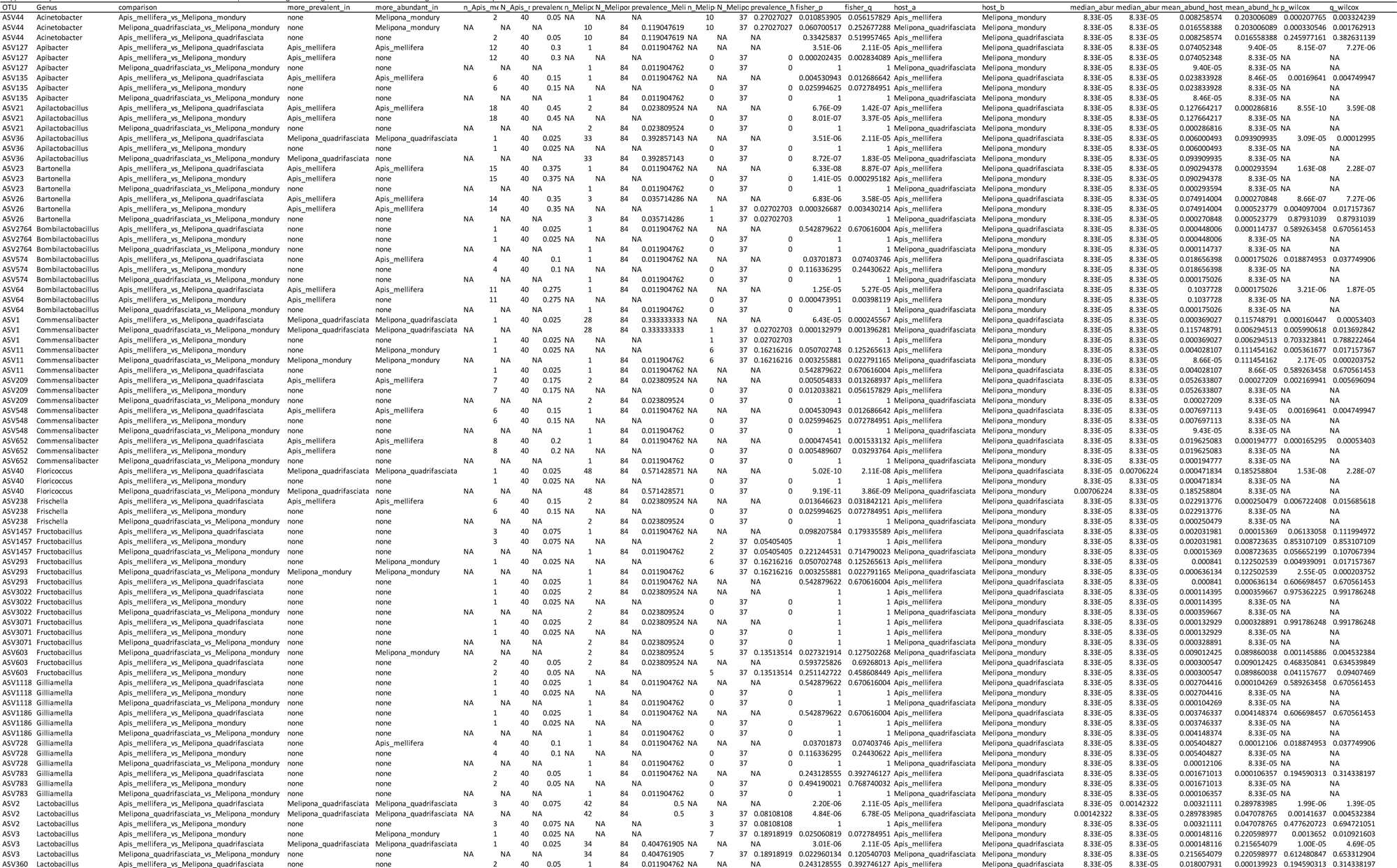
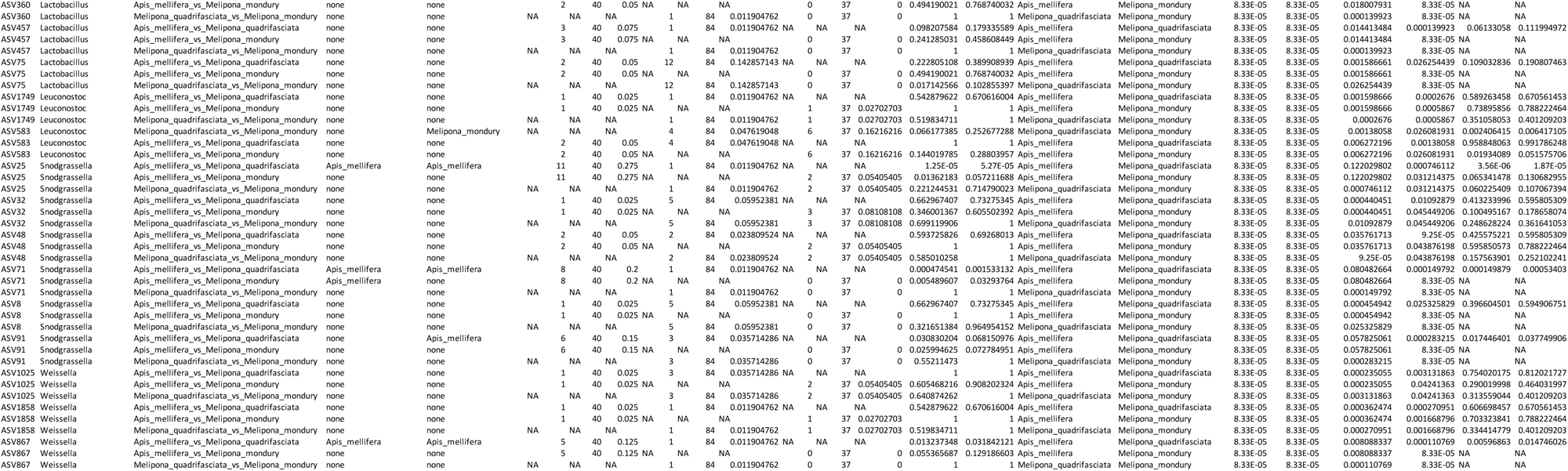
Prevalence and relative abundance statistic comparisons.

**Supplementary File 4.**
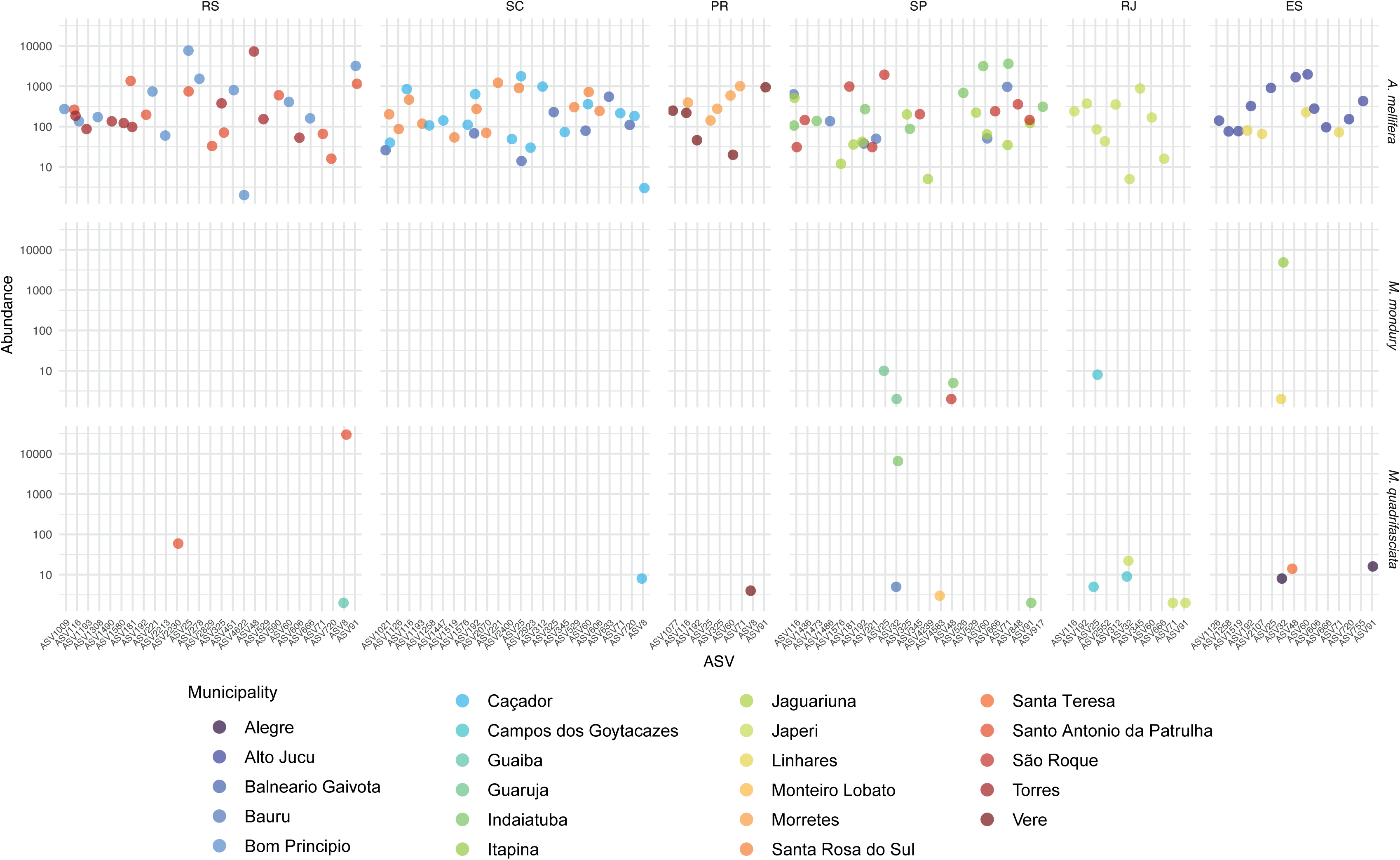
Total number of reads of *Snodgrassella* ASVs per sample

**Supplementary File 5.**
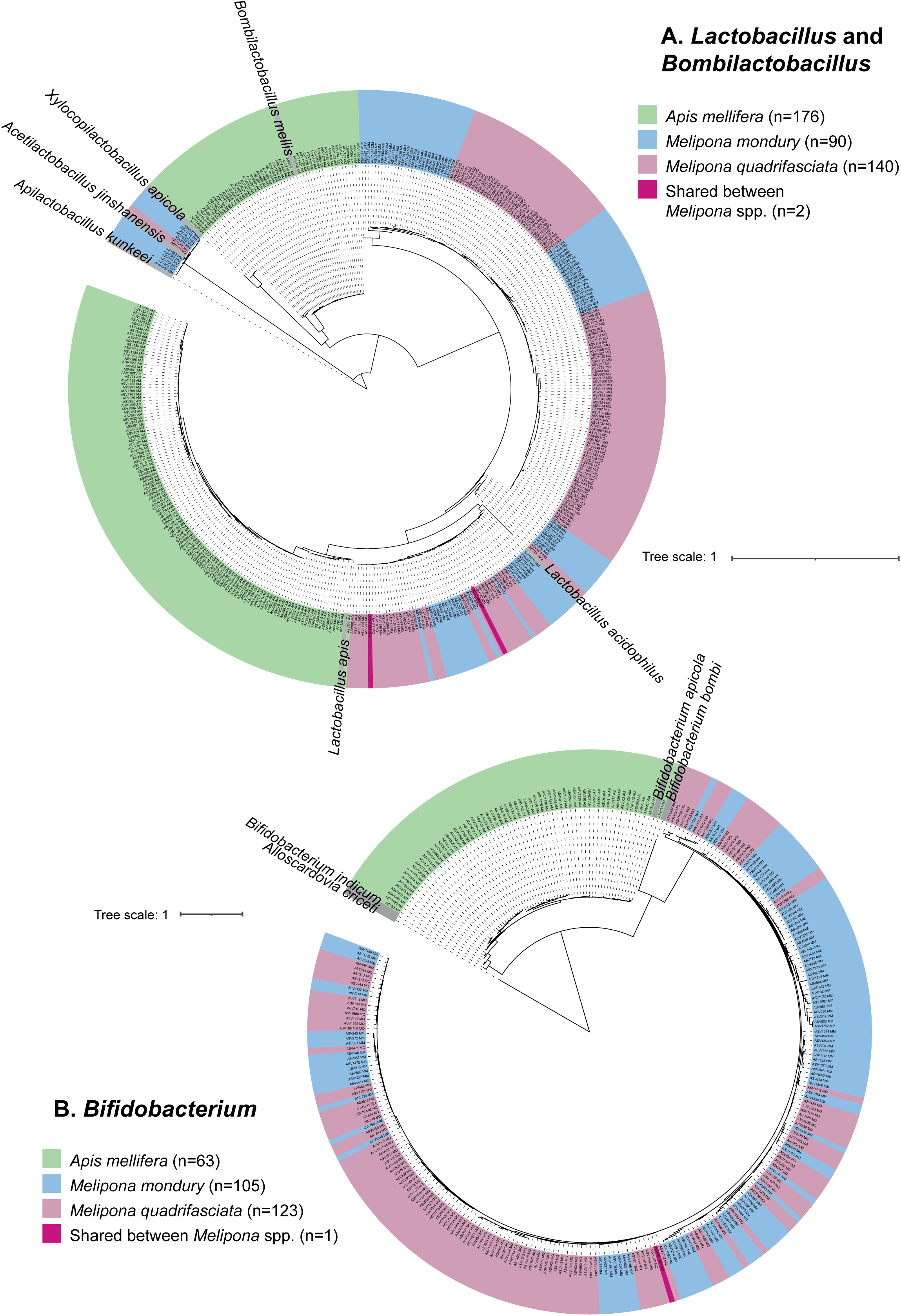
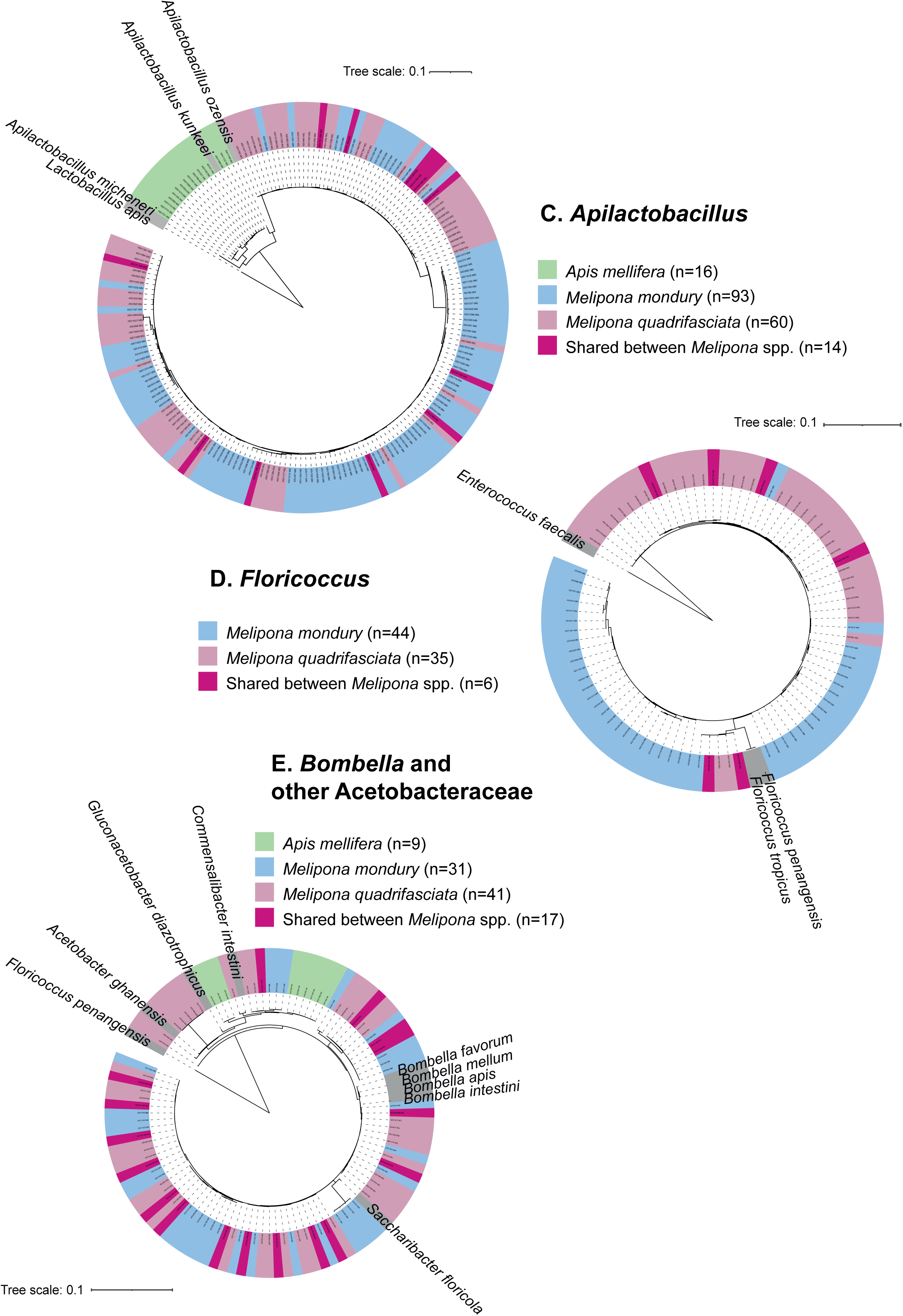
Phylogenetic trees of ASVs from the “common core” bacteria.

## References

1. Martinson VG, Moy J, Moran NA. Establishment of characteristic gut bacteria during development of the honeybee worker. Appl Environ Microbiol 2012; 78:2830–2840. 10.1128/AEM.07810-11

2. Powell JE et al. Routes of acquisition of the gut microbiota of the honey bee *Apis mellifera*. Appl Environ Microbiol 2014; 80:7378–7387. 10.1128/AEM.01861-14

3. Kešnerová L et al. Disentangling metabolic functions of bacteria in the honey bee gut. PLoS Biol 2017; 15:e2003467. 10.1371/journal.pbio.2003467

4. Ellegaard KM et al. Genomic changes underlying host specialization in the bee gut symbiont *Lactobacillus* Firm5. Mol Ecol 2019; 28:2224–2237. 10.1111/mec.15075

5. Motta EVS et al. Host-microbiome metabolism of a plant toxin in bees. eLife 2022; 11:e82595. 10.7554/eLife.82595

6. Zhang Z et al. Honeybee gut *Lactobacillus* modulates host learning and memory behaviors via regulating tryptophan metabolism. Nat Commun 2022; 13:2037. 10.1038/s41467-022-29760-0

7. Li L et al. Gut microbiome drives individual memory variation in bumblebees. Nat Commun 2021; 12:6588. 10.1038/s41467-021-26833-4

8. Koch H, Schmid-Hempel P. Socially transmitted gut microbiota protect bumble bees against an intestinal parasite. Proc Natl Acad Sci U S A (PNAS) 2011; 108:19288–19292. 10.1073/pnas.1110474108

9. Raymann K, Shaffer Z, Moran NA. Antibiotic exposure perturbs the gut microbiota and elevates mortality in honeybees. PLoS Biol 2017; 15:e2001861. 10.1371/journal.pbio.2001861

10. Motta EVS, Moran NA. The honeybee microbiota and its impact on health and disease. Nat Rev Microbiol 2024; 22:122–137. 10.1038/s41579-023-00990-3

11. Engel P et al. The Bee Microbiome: Impact on bee health and model for evolution and ecology of host-microbe interactions. mBio 2016; 7:e02164–15. 10.1128/mBio.02164-15

12. Engel P, Martinson VG, Moran NA. Functional diversity within the simple gut microbiota of the honey bee. Proc Natl Acad Sci U S A (PNAS) 2012; 201202970. 10.1073/pnas.1202970109

13. Zhong Z et al. Gut symbiont-derived anandamide promotes reward learning in honeybees by activating the endocannabinoid pathway. Cell Host Microbe 2024; 32:1944–1958.e7. 10.1016/j.chom.2024.09.013

14. Cabirol A et al. A defined community of core gut microbiota members promotes cognitive performance in honey bees. bioRxiv 2023; 2023.01.03.522593. 10.1101/2023.01.03.522593

15. Kwong WK et al. Dynamic microbiome evolution in social bees. Sci Adv 2017; 3:e1600513. 10.1126/sciadv.1600513

16. Prasad A et al. Evolution of gut microbiota across honeybee species revealed by comparative metagenomics. Nat Commun 2025; 16:9069. 10.1038/s41467-025-64115-5

17. Sarton-Lohéac G et al. Deep divergence and genomic diversification of gut symbionts of neotropical stingless bees. mBio 2023; 14:e03538–22. 10.1128/mbio.03538-22

18. Leonhardt SD, Kaltenpoth M. Microbial communities of three sympatric Australian stingless bee species. PLoS One 2014; 9:e105718. 10.1371/journal.pone.0105718

19. Cerqueira AES et al. *Melipona* stingless bees and honey microbiota reveal the diversity, composition, and modes of symbionts transmission. FEMS Microbiol Ecol 2024; 100:fiae063. 10.1093/femsec/fiae063

20. Haag KL et al. Temporal changes in gut microbiota composition and pollen diet associated with colony weakness of a stingless bee. Microb Ecol 2023; 85:1514– 1526. 10.1007/s00248-022-02027-3

21. Roffet-Salque M et al. Widespread exploitation of the honeybee by early Neolithic farmers. Nature 2016; 534:S17–S18. 10.1038/nature18451

22. Dunne J et al. Honey-collecting in prehistoric West Africa from 3500 years ago. Nat Commun 2021; 12:2227. 10.1038/s41467-021-22425-4

23. Paris EH et al. The origins of Maya stingless beekeeping. J Ethnobiol 2020; 40:386–405. 10.2993/0278-0771-40.3.386

24. Cerqueira AES et al. Extinction of anciently associated gut bacterial symbionts in a clade of stingless bees. ISME J 2021; 15:2813–2816. 10.1038/s41396-021-01000-1

25. Kueneman JG et al. Neotropical bee microbiomes point to a fragmented social core and strong species-level effects. Microbiome 2023; 11:150. 10.1186/s40168-023-01593-z

26. Ramírez-Ahuja M de L, et al. Gut microbiota diversity in 16 stingless bee species (Hymenoptera: Apidae: Meliponini). Microorganisms 2025; 13:1645. 10.3390/microorganisms13071645

27. Batalha-Filho H et al. Phylogeography and historical demography of the neotropical stingless bee *Melipona quadrifasciata* (Hymenoptera, Apidae): incongruence between morphology and mitochondrial DNA. Apidologie 2010; 41:14. 10.1051/apido/2010001

28. Batalha-Filho H et al. Geographic distribution and spatial differentiation in the color pattern of abdominal stripes of the neotropical stingless bee *Melipona quadrifasciata* (Hymenoptera: Apidae). Zoologia 2009; 26:213–219. 10.1590/S1984-46702009000200003

29. Françoso E, Arias MC. Cytochrome c oxidase I primers for corbiculate bees: DNA barcode and mini-barcode. Mol Ecol Resour 2013; 13:844–850. 10.1111/1755-0998.12135

30. Simon C et al. Evolution, weighting, and phylogenetic utility of mitochondrial gene sequences and a compilation of conserved Polymerase Chain Reaction primers. Ann Entomol Soc Am 1994; 87:651–701. 10.1093/aesa/87.6.651

31. Edgar RC. MUSCLE: multiple sequence alignment with high accuracy and high throughput. Nucleic Acids Res 2004; 32:1792–1797. 10.1093/nar/gkh340

32. Clement M, Posada D, Crandall KA. TCS: a computer program to estimate gene genealogies. Mol Ecol 2000; 9:1657–1659. 10.1046/j.1365-294x.2000.01020.x

33. Callahan BJ et al. DADA2: High-resolution sample inference from Illumina amplicon data. Nat Methods 2016; 13:581–583. 10.1038/nmeth.3869

34. Quast C et al. The SILVA ribosomal RNA gene database project: improved data processing and web-based tools. Nucleic Acids Res 2013; 41:D590–596. 10.1093/nar/gks1219

35. Wang Q et al. Naïve bayesian classifier for rapid assignment of rRNA sequences into the new bacterial taxonomy. Appl Environ Microbiol 2007; 73:5261–5267. 10.1128/AEM.00062-07

36. Guindon S et al. New algorithms and methods to estimate maximum-likelihood phylogenies: Assessing the performance of PhyML 3.0. Syst Biol 2010; 59:307–321. 10.1093/sysbio/syq010

37. R Core Team. R: The R project for statistical computing. https://www.r-project.org/. (2025, date last accessed).

38. McMurdie PJ, Holmes S. phyloseq: An R package for reproducible interactive analysis and graphics of microbiome census data. PLoS One 2013; 8:e61217. 10.1371/journal.pone.0061217

39. Neu AT, Allen EE, Roy K. Defining and quantifying the core microbiome: Challenges and prospects. Proc Natl Acad Sci U S A (PNAS) 2021; 118:e2104429118. 10.1073/pnas.2104429118

40. Oksanen J et al. vegan: Community Ecology Package. https://CRAN.R-project.org/package=vegan. (2025, date last accessed).

41. Kelly BJ et al. Power and sample-size estimation for microbiome studies using pairwise distances and PERMANOVA. Bioinformatics 2015; 31:2461–2468. 10.1093/bioinformatics/btv183

42. Lin H, Peddada SD. Multigroup analysis of compositions of microbiomes with covariate adjustments and repeated measures. Nat Methods 2024; 21:83–91. 10.1038/s41592-023-02092-7

43. Peschel S et al. NetCoMi: network construction and comparison for microbiome data in R. Brief Bioinform 2021; 22:bbaa290. 10.1093/bib/bbaa290

44. Kwong WK, Moran NA. Cultivation and characterization of the gut symbionts of honey bees and bumble bees: description of *Snodgrassella alvi* gen. nov., sp. nov., a member of the family Neisseriaceae of the Betaproteobacteria, and *Gilliamella apicola* gen. nov., sp. nov., a member of Orbaceae fam. nov., Orbales ord. nov., a sister taxon to the order ‘Enterobacteriales’ of the Gammaproteobacteria. Int J Syst Evol Microbiol 2013; 63:2008–2018. 10.1099/ijs.0.044875-0

45. Santini AT et al. Gut microbiota of Brazilian *Melipona* stingless bees: dominant members and their localization in different gut regions. bioRxiv 2025., 2025.06.03.657762. 10.1101/2025.06.03.657762

46. Díaz S et al. Report on the microbiota of *Melipona quadrifasciata* affected by a recurrent disease. J Invertebr Pathol 2017; 143:35–39. 10.1016/j.jip.2016.11.012

47. Luo S, Zhang X, Zhou X. Temporospatial dynamics and host specificity of honeybee gut bacteria. Cell Rep 2024; 43:114408. 10.1016/j.celrep.2024.114408

48. Liu H et al. Gut microbial diversity in stingless bees is linked to host wing size and is influenced by geography. J Invertebr Pathol 2023; 198:107909. 10.1016/j.jip.2023.107909.

49. Mills TJT et al. Hive transplantation has minimal impact on the core gut microbiome of the Australian stingless bee, *Tetragonula carbonaria*. Microb Ecol 2023; 86:2086–2096. 10.1007/s00248-023-02222-w

50. Hall MA et al. Temporal changes in the microbiome of stingless bee foragers following colony relocation. FEMS Microbiol Ecol 2021; 97. 10.1093/femsec/fiaa236

